# Ranking quantitative resistance to Septoria tritici blotch in elite wheat cultivars using automated image analysis

**DOI:** 10.1101/129353

**Authors:** Petteri Karisto, Andreas Hund, Kang Yu, Jonas Anderegg, Achim Walter, Fabio Mascher, Bruce A. McDonald, Alexey Mikaberidze

## Abstract

Quantitative resistance is likely to be more durable than major gene resistance for controlling Septoria tritici blotch (STB) on wheat. Earlier studies hypothesized that resistance affecting the degree of host damage, as measured by the percentage of leaf area covered by STB lesions, is distinct from resistance that affects pathogen reproduction, as measured by the density of pycnidia produced within lesions. We tested this hypothesis using a collection of 335 elite European winter wheat cultivars that was naturally infected by a diverse population of *Zymoseptoria tritici* in a replicated field experiment. We used automated image analysis (AIA) of 21420 scanned wheat leaves to obtain quantitative measures of conditional STB intensity that were precise, objective, and reproducible. These measures allowed us to explicitly separate resistance affecting host damage from resistance affecting pathogen reproduction, enabling us to confirm that these resistance traits are largely independent. The cultivar rankings based on host damage were different from the rankings based on pathogen reproduction, indicating that the two forms of resistance should be considered separately in breeding programs aiming to increase STB resistance. We hypothesize that these different forms of resistance are under separate genetic control, enabling them to be recombined to form new cultivars that are highly resistant to STB. We found a significant correlation between rankings based on automated image analysis and rankings based on traditional visual scoring, suggesting that image analysis can complement conventional measurements of STB resistance, based largely on host damage, while enabling a much more precise measure of pathogen reproduction. We showed that measures of pathogen reproduction early in the growing season were the best predictors of host damage late in the growing season, illustrating the importance of breeding for resistance that reduces pathogen reproduction in order to minimize yield losses caused by STB. These data can already be used by breeding programs to choose wheat cultivars that are broadly resistant to naturally diverse *Z. tritici* populations according to the different classes of resistance.

---

*Zymoseptoria tritici* (Desm.) Quaedvlieg & Crous (formerly *Mycosphaerella graminicola* (Fuckel) J. Schrot. in Cohn) is a fungal pathogen that poses a major threat to wheat production, particularly in temperate areas (Jørgensen et al., 2014; Dean et al., 2012). It infects wheat leaves, causing the disease Septoria tritici blotch (STB). Yield losses caused by STB can be 5-10%, even when resistant cultivars and fungicides are used in combination. Around 1.2 billion dollars are spent annually in Europe on fungicides targeted mainly towards STB control (Torriani et al., 2015). *Z. tritici* populations in Europe carry a high degree of fungicide resistance (reviewed in Fones and Gurr, 2015; Torriani et al., 2015). In several cases, fungicides repeatedly lost their efficacy only a few years after their introduction due to rapid emergence of fungicide-resistant strains of *Z. tritici* (Griffin and Fisher, 1985; Fraaije et al., 2005; Torriani et al., 2009). Resistance to azoles, an important class of fungicides that is widely used to control STB, has been growing steadily over the last twenty years in Europe (Cools and Fraaije, 2013; Zhan et al., 2006) and appeared recently in North America (Estep et al., 2015). Therefore, STB-resistant wheat cultivars have become an important breeding objective to enable more effective management of the disease (McDonald and Mundt, 2016). Major resistance genes such as *Stb6* (Brading et al., 2002) provide nearly complete resistance against a subset of *Z. tritici* strains carrying the wild type *AvrStb6* allele (Zhong et al., 2017), but as found for fungicides, major resistance often breaks down a few years after it is introduced (Cowger et al., 2000). Quantitative resistance may be conferred by a large number of quantitative trait loci (QTLs) with small and additive effects that can be combined to provide high levels of disease resistance (Poland et al., 2009; St. Clair, 2010; Kou and Wang, 2010; McDonald and Linde, 2002; Mundt, 2014). Quantitative resistance is thought to be more durable and hence deserves more attention from breeders (McDonald and Linde, 2002; St. Clair, 2010; Mundt, 2014).

To enable breeding for quantitative resistance to STB, we need to comprehensively analyze the quantitative distribution of its associated phenotypes, which is much more difficult than phenotyping major gene resistance that typically shows a binomial distribution. This challenge was recognized more than forty years ago and a number of studies were conducted to evaluate quantitative resistance to STB under field conditions using artificial inoculation (Rosielle, 1972; Shaner and Finney, 1982; Eyal, 1992; Brown et al., 2001; Miedaner et al., 2013) and natural infection (Rosielle, 1972; Shaner et al., 1975; Miedaner et al., 2013; Kollers et al., 2013*b*). Resistance to STB was also investigated on detached leaves with artificial inoculations (e. g. by Chartrain et al., 2004). Several studies performed visual scoring of quantitative resistance only once during the growing season (Rosielle, 1972; Shaner and Finney, 1982; Eyal, 1992; Miedaner et al., 2013), while other studies included two or more time points (Shaner et al., 1975; Brown et al., 2001; Kollers et al., 2013*b*). One of the most comprehensive early studies screened 7500 wheat varieties including 2000 durum wheat cultivars to select the 460 most resistant varieties for more detailed visual scoring (Rosielle, 1972).

Understanding the infection cycle of STB in important to distinguish and measure the most relevant aspects of quantitative resistance to the disease. *Z. tritici* spores germinate on wheat leaves and penetrate the leaves through stomata (Kema et al., 1996). After penetration, the fungus grows for several days within leaves without producing visible symptoms. During this latent phase, *Z. tritici* mycelium grows in the apoplast and invades the host mesophyll around the position of the initial penetration (Duncan and Howard, 2000). After 10-14 days of asymptomatic growth, the fungus becomes necrotrophic, necrotic lesions appear in the invaded host tissue and asexual fruiting bodies called pycnidia begin to form (Kema et al., 1996; Duncan and Howard, 2000). In the dead host tissue, the fungus continues to grow saprotrophically and produces sexual fruiting bodies called pseudothecia 25-30 days after infection (Sanchez-Vallet et al., 2015). Whether *Z. tritici* is best referred to as a hemibiotroph or a latent necrotroph remains unclear (Sanchez-Vallet et al., 2015). Asexual pycnidiospores are usually spread by rain splash while sexual ascospores are spread by wind. The pathogen typically undergoes 5-6 rounds of asexual reproduction and 1-2 rounds of sexual reproduction per growing season.

Studies by Zhan et al. (1998) and Zhan et al. (2000) indicated that *≈*66% of infections on flag leaves came from asexual spores, while *≈*24% came from ascospores originating from within the infected field and *≈*10% of flag leaf infections were immigrants from surrounding fields. Pathogen asexual reproduction is thus the most important factor explaining infection on flag leaves, but a significant fraction of flag leaf infections can originate from airborne ascospores coming from within or outside of the field. The amount of necrosis induced by STB on the uppermost leaves determines yield losses (Brokenshire, 1976).

The amount of STB can be measured by determining disease incidence, disease severity and pycnidia density (Madden et al., 2007; Shaner and Finney, 1982). Disease incidence is the proportion of plant units diseased (Madden et al., 2007). In the case of STB, the relevant plant units are leaves, hence we consider STB incidence to be the proportion of wheat leaves diseased. The degree of infection on a leaf is a measure of disease severity. STB severity is typically measured as the percentage of leaf area covered by lesions (PLACL). Mean STB severity in a plot is usually defined as an average value across a random sample that includes both infected and uninfected leaves. In contrast, conditional STB severity is defined as the mean severity in a sample that includes only infected leaves. Following Rosielle (1972), Shaner et al. (1975) and Shaner and Finney (1982), we consider pycnidia density as another important measure of STB. We use the term “disease intensity” as a general term that refers to both disease severity and pycnidia density. Similar to the definition of conditional severity, we define conditional pycnidia density as the average pycnidia density in a sample of infected leaves. For this study, we collected and analyzed only infected leaves and therefore we did not measure STB incidence. The measures of STB that we report here represent conditional severity and conditional pycnidia density.

Earlier studies of STB resistance combined disease severity and incidence using visual assessments based on categorical scales. In studies of Rosielle (1972), Shaner et al. (1975) and Eyal (1992) these scales included both the degree of lesion coverage and the density of pycnidia in lesions, but in the studies of Brown et al. (2001) and Chartrain et al. (2004) the disease scores were based on leaf coverage by lesions bearing pycnidia (i. e. using a presence/absence measurement of pycnidia to define STB lesions). The accuracy of all these methods is limited by an inherent subjective bias and a small number of qualitative categories, which may limit success in breeding for STB resistance.

In several studies, the importance of resistance that suppresses pathogen reproduction (i. e. production of pycnidia) was recognized based on qualitative observations of pycnidia coverage (e. g. Rosielle, 1972; Shaner et al., 1975; Shaner and Finney, 1982). Because manual counting of pycnidia is extremely labor-intensive, it was only feasible to count pycnidia on a very small scale (e. g. Shaner et al., 1975)] before the development of new technology to automate this process.

Automated image analysis (AIA) provides a promising tool for measuring quantitative disease resistance in the field (Mahlein, 2016; Simko et al., 2017). Mutka and Bart (2015) and Mahlein (2016) highlight the importance of standardized imaging methods for reproducibility. We used a novel phenotyping method based on automated analysis of scanned leaf images (Stewart and McDonald, 2014; Stewart et al., 2016) in a winter wheat panel planted to 335 elite European cultivars in a replicated field experiment (Kirchgessner et al., 2017). This method benefits from well defined procedure of detaching leaves and scanning them under standardized conditions, thus leading to objective and reproducible results. Additionally, it enables generation of large amounts of reliable data at a relatively low cost.

Importantly, our AIA method allowed us to quantitatively separate the net effects of resistance components affecting host damage from resistance components affecting pathogen reproduction. Components of resistance (Parlevliet, 1979) are defined as resistance factors suppressing individual processes of the infection cycle (Willocquet et al., 2017). Components of resistance that suppress infection efficiency and lesion expansion are responsible for a reduction in host damage, while components of resistance that suppress spore production (Parlevliet, 1979) or pycnidia coverage (Simon and Cordo, 1998) are responsible for a reduction in pathogen reproduction Pathogen reproduction was quantified by automatic counting of asexual fruiting bodies of the pathogen (pycnidia) on wheat leaves (Stewart and McDonald, 2014; Stewart et al., 2016) and host damage was measured by automatic detection of discolored leaf areas caused by STB infection.

In this large-scale field experiment, leaves were infected naturally by a genetically diverse local population of *Z. tritici* and the epidemic was allowed to develop naturally. Despite three fungicide treatments including five active ingredients that eliminated virtually all other diseases, STB infection was widespread across the field experiment. This pervasive natural infection by a fungicide-resistant population allowed us to investigate quantitative resistance in a nearly pure culture of *Z. tritici* under the high-input field conditions typical of Europe. The combination of wet and cool weather conditions favoring development of STB, a large number of wheat cultivars planted in a single location, and utilization of a novel AIA method enabled a comprehensive characterization of quantitative resistance that led to a clear ranking of STB resistance in a broad collection of European winter wheat cultivars.

We report separate rankings of wheat cultivars based on two different components of epidemic outcome, one measured as host damage and the other as pathogen reproduction. We found that the two rankings are considerably different. We identified a phenotypic quantity that combines these two components and found that it correlates best with the ranking based on traditional visual assessments. In this way, we identified new, broadly active sources of resistance to STB in existing European wheat cultivars. Our findings open several possibilities for further genetic studies of quantitative resistance to STB.

## Materials and Methods

### Plant materials and experimental design

A total of 335 elite European winter wheat (*Triticum aestivum*) varieties from the GABI-wheat panel (Kollers et al., 2013*a*,*b*) were evaluated in this experiment. Two biological replicates of the wheat panel were grown during the 2015–2016 growing season in two complete blocks separated by about 100 meters at the Field Phenotyping Platform (FIP) site of the Eschikon Field Station of the ETH Zurich, Switzerland (coordinates 47.449, 8.682) (Kirchgessner et al., 2017). The complete blocks were composed of 18 rows and 20 columns composed of 1.2m by 1.7m plots, with the genotypes arranged randomly within each block except for a check variety (CH Claro) that was planted at least once within each row and column, leading to 21 replicates of CH Claro within each block. All plots were sown on 13 October 2015.

including Practices recommended for conventional, high-input wheat production in Switzerland include applications of fertilizers and pesticides. Fertilizers were applied five times during spring 2016, including boron with ammonium nitrate (nitrogen 52 kg/ha) on 4 March; P_2_O_5_ at 92 kg/ha on 7 March; K_2_O at 120 kg/ha on 10 March; magnesium with ammonium nitrate on 12 April (magnesium 15 kg/ha, nitrogen 72 kg/ha) and on 20 May (magnesium 4 kg/ha, nitrogen 19 kg/ha). The pre-emergence herbicide Herold SC (Bayer) was applied on 29 October 2015 (dose 0.6 l/ha). The stem shortener Moddus (Syngenta) was applied on 6 April 2016 (dose 0.4 l/ha) at GS (growth stage) 31 that corresponds to stem elongation (Zadoks et al., 1974). Insecticide Biscaya (Bayer) was applied on 25 May 2016 (dose 0.3 l/ha) at GS 51 that corresponds to inflorescence emergence.

Fungicides were applied three times: (i) 6 April, Input, Bayer (a mixture of spiroxamin at 300 g/l and prothioconazole at 150 g/l, dose 1.25 l/ha, GS 31); (ii) 25 May, Aviator Xpro, Bayer (a mixture of bixafen at 75 g/l and prothioconazole at 150 g/l, dose 1.25 l/ha, GS 51) and 6 June, Osiris, BASF (a mixture of epoxiconazole at 56.25 g/l and metconazole at 41.25g/l, dose 2.5 l/ha, GS 65 that corresponds to anthesis). In total, the three fungicide applications included five active ingredients representing three modes of action. In particular, bixafen is a succinate dehydrogenase inhibitor (SDHI); prothioconazole, epoxiconazole and metconazole are azoles, the main chemical group in the sterol 14a-demethylation inhibitors (DMI) class of fungicides; and spiroxamin belongs to the morpholine class of fungicides. This strategy of fungicide application aimed to minimize the overall levels of the most common fungal diseases on Swiss wheat.

### STB inoculum and calculation of number of cycles of infection

All STB infection was natural, with the majority of primary inoculum likely originating from airborne ascospores coming from nearby wheat fields that surround the Eschikon field site. We obtained rough estimates of the average number of asexual cycles of reproduction within the pathogen population. For this purpose, we used the data from Shaw (1990) showing the effect of temperature on latent period and local weather data coming from the nearby Lindau weather station (see Appendix A.1 for details of estimation).

### Disease assessment based on automated image analysis

Leaves exhibiting obvious STB lesions were collected two times during the growing season. The first collection was made on 20 May 2016 (*t*_1_, approximately GS 41 that corresponds to booting) and the second collection was made on 4 July 2016 (*t*_2_, approximate GS was in the range 75-85 that corresponds to milk development (75) and dough development (85)). For both collections, 16 infected leaves were collected at random from each plot. At *t*_1_, leaves were collected from the highest infected leaf layer, which was typically the third or fourth fully extended, but non-senescent leaf still visible when counting from the ground or one to three leaf layers below the top leaf. At *t*_2_, the leaf layer below the flag leaf (F-1) was sampled in each plot. The sampled leaves were placed in paper envelopes, kept on ice in the field, and stored at 4° C for two days before mounting on A4 paper with printed reference marks and sample names, as described in (Stewart et al., 2016). Absorbent paper was placed between each sheet of eight mounted leaves and sheets were pressed with approximately 5 kg at 4° C for two-three days prior to scanning at 1200 dpi with a Canon CanoScan LiDE 220 flatbed scanner. The resulting scans were saved as “jpeg” images.

Scanned images were analyzed with the software ImageJ (Schindelin et al., 2015) using a modification of the macro described by Stewart and McDonald (2014) and Stewart et al. (2016)(source code of the macro and a user manual are given in (Stewart et al., 2016)). The parameters used for the macro are given in Supplemental Table S1 and an explanation of their meaning is provided in the macro instructions. Figure 1 illustrates the workflow associated with the macro. The maximum length of the scanned area for each leaf was 17 cm. When leaves were longer than 17 cm, bases of the leaves were placed within the scanned area, while the leaf tips extended outside the scanned area. For each leaf, the following quantities were automatically recorded from the scanned image: total leaf area, necrotic and chlorotic leaf area, number of pycnidia and their positions on the leaf. Necrotic and chlorotic leaf areas were detected based on discoloration of the leaf surface and were not based on the presence of pycnidia. From these measurements, we calulated the percentage of leaf area covered by lesions (PLACL), the density of pycnidia per unit lesion area (*ρ*_lesion_), the density of pycnidia per unit leaf area (*ρ*_leaf_) (see Table 1).

**Table 1:**
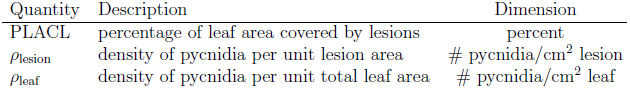
Important STB disease properties determined using automated image analysis.

**Figure 1:**
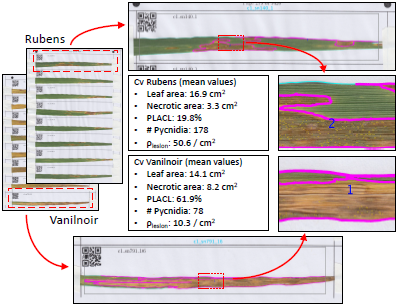
Illustration of the automated phenotyping procedure with the ImageJ macro (Stewart et al., 2016). Leaves are mounted on pre-printed paper sheets; the ImageJ macro distinguishes leaves from the white background; within each leaf, the macro identifies necrotic lesions (lesion labels are shown in blue) and their areas; within each lesion, the macro identifies fungal fruiting bodies called pycnidia (black dots).

The three quantities PLACL, *ρ*_lesion_, and *ρ*_leaf_ quantify different aspects of conditional intensity of STB in each plot. Although we aimed to collect only infected leaves, there were a few cases in plots with very little STB when the collected leaves did not have necrotic lesions or did not have pycnidia. These leaves were not considered when calculating the mean or median values used for ranking cultivars or assessing the magnitude of effects.

We identified several cases of biased leaf collection at *t*_1_, where leaves were sampled from lower leaf layers in which some of the leaves exhibited natural senescence, leading to extensive chlorosis and some necrosis. We removed several sheets that were strongly affected by collector bias or natural senescence. Next, we estimated the proportion of remaining scoring errors due to collector bias as *p*_cb_ = 0.055 to extensive chlorosis and some necrosis. We removed several sheets that were strongly affected by collector bias or natural senescence. Next, we estimated the proportion of remaining scoring errors due to collector bias as *p*_cb_ = 0.055 *±* 0.003 by testing 200 leaves sampled randomly from the entire leaf population.

We found that the original image analysis macro reported in Stewart et al. (2016) identified many falsely detected pycnidia, especially in cultivars with large numbers of thick trichomes. We addressed this issue by improving the macro and performing an extensive optimization of the macro parameters. We changed the macro code to improve detection of pycnidia by enhancing the contrast and optimizing the proportion of red, green and blue channels in the leaf images. To find an optimal set of macro parameters, we tested more than 160 different sets of parameters on 16 leaves that represented a wide range of numbers of pycnidia per leaf. We chose the parameter set that had the highest concordance coefficient between the numbers of pycnidia per leaf detected by the macro and the numbers of pycnidia counted manually. We then analyzed the entire set of images using this optimized parameter set and tested the outcome in several ways.

First we visually inspected 209 leaves chosen using random stratified sampling within the range *ρ*_lesion_ *<* 50. We focused on this range, because we found it to be most prone to false detection of pycnidia. For each leaf subjected to visual inspection, we performed a binary (yes/no) assessment of whether the macro correctly detected most of the lesions and pycnidia. This was done by comparing scanned leaf images with overlay images in which the detected lesions and pycnidia were marked by the macro (examples of overlays are shown in Fig. 1). Errors that led to incorrect quantification of disease symptoms could be divided into four categories: defects on leaves, collector bias, scanning errors and deficiencies in the image analysis macro. Leaf defects included insect damage, mechanical damage, insect bodies and frass, other fungi, uneven leaf surfaces creating shadows, and dust particles on leaves. Scanning errors included shadows on leaf edges and folded leaves that resulted from leaves that were not properly flattened prior to scanning. Deficiencies of the macro consisted of recognizing green parts of leaves as lesions, or recognizing dark spaces between light-colored leaf hairs as pycnidia, and parts of dark borders around lesions as pycnidia. In total, 21 leaves were deemed to exhibit scoring errors and were removed from the dataset as a result of this procedure. Based on this outcome, we estimated the proportion of scoring errors as *p*_tot_ = 0.10 *±* 0.02 out of which proportion of errors due to false pycnidia was estimated as *p*_fp_ = 0.004 *±* 0.03. Here, the uncertainties are reported in the form of the 95 % confidence intervals calculated according to *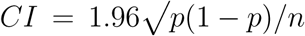*, where *p* is the sample proportion and *n* is the sample size.

To further quantify the accuracy of pycnidia detection by the macro, we compared pycnidia counts by the macro to manual pycnidia counts. For this purpose, we selected 30 random leaves from the entire dataset as well as 6 leaves with more than 400 counted pycnidia. We found that pycnidia counts by the macro had high concordance (concordance coefficient *r*_*c*_ = 0.97) and high correlation (*r*_*s*_ = 0.96, *p* = 5 10^17^) with manual pycnidia counts. The macro underestimated pycnidia counts on average by 26 *±* 10% (the uncertainty here represents the 95% confidence interval).

### Disease assessment based on visual scoring

Visual assessments of STB were performed at three time points: 20 May (approximately GS 41), 21 June (approximately GS 75) and 29 June (approximately GS 80). The STB level in each plot was scored by global assessment of the three uppermost leaf layers on a 1-9 scale (1 means no disease, 9 means complete infection, please refer to Supplemental Table S2 for more details of the scale) based on both STB incidence and severity (Michel, 2001). The presence of pycnidia was used as an indicator of STB infection. The absence of pycnidia was interpreted as an absence of STB, even if necrotic lesions were visible. During visual scoring, the presence of other diseases (such as stripe rust, septoria nodorum blotch and fusarium head blight) was assessed qualitatively. All plots were scored with approximately equal time spent on each plot.

### Statistical analysis

We compared differences in STB resistance among cultivars for each dataset by pooling together the data points from individual leaves belonging to different replicates and sampling dates. The data from the automated image analysis consisted of *≈* 60 data points per cultivar representing the two time points and the two biological replicates (blocks). Cultivar CH Claro was an exception because it was replicated 42 times and thus had *≈* 1300 data points from leaf image analysis and 124 data points from visual scoring. The relative STB resistance of all wheat cultivars was ranked based on the means of the conditional measurements of disease intensity PLACL, *ρ*_lesion_ and *ρ*_leaf_ over *≈* 60 individual leaf data points. We also calculated medians and standard errors of the means for each cultivar.

Resistance that affects STB incidence was not measured using AIA, as only diseased leaves were analyzed. The visual scoring data was based on three time points and two biological replicates, generating 6 data points per cultivar. For each cultivar the area under disease progress curve (AUDPC) was calculated by taking averages of visual scores over the two replicates. It was assumed that infection started from zero at 14 days before the first assessment. To analyze differences between cultivars these scores were weighted with coefficients that depend on times of assessments such that each weighted score gives a proportional contribution to the total AUDPC and the average over scores from different replicates and time points gives the total AUDPC (see Appendix A.2 for details on calculation of AUDPC and weighting of scores). AUDPC was used to rank the cultivars according to visual scoring (Fig. 3D).

The significance of differences in resistance between cultivars was tested with the global Kruskal-Wallis test (Sokal and Rohlf, 2012) using the kruskal.test function in R (R Core Team, 2016) for each data set. For resistance measures showing global differences between cultivars, we determined significantly different groups of cultivars based on pairwise comparisons, whereby any two cultivars in the same group were not significantly different from each other and any two cultivars from different groups were different. Pairwise differences were tested with multiple pairwise Kruskal-Wallis tests using the function kruskal in the package agricolae in R (de Mendiburu, 2016) using the false discovery rate (FDR) 0.05 for significance level correction (Benjamini and Hochberg, 1995) for multiple comparisons.

We determined correlations between cultivar rankings based on AUDPC and using both means and medians of PLACL, *ρ*_lesion_ and *ρ*_leaf_ for each cultivar. We also computed correlations with respect to means of PLACL, *ρ*_lesion_ and *ρ*_leaf_ between *t*_1_ and *t*_2_ to determine the predictive power of these quantities. In this case, means were taken over about 15 leaves originating from the same plot. All correlations were calculated with Spearman’s correlation test (Sokal and Rohlf, 2012) using the open-source scipy package (http://www.scipy.org) written for the Python programming language (http://www.python.org).

We analyzed differences between *t*_1_ and *t*_2_ in terms of PLACL, *ρ*_lesion_ and *ρ*_leaf_ to identify cultivars whose resistance increased over time. For this purpose, we used the Wilcoxon rank sum test with the FDR correction for multiple comparisons at *p <* 0.01.

### Workflow

For clarity, we briefly summarize the workflow of the experiment as a linear sequence of steps. We first collected leaf samples, mounted them on sheets of paper and scanned them. Next, we performed the basic image analysis using the ImageJ macro as illustrated in Fig. 1. After that, we tested the data and estimated scoring errors using visual examination of leaf images. Finally, we performed detailed statistical analyses using the verified datasets.

## Results

### Overall description of the STB epidemic

Despite three fungicide applications with five active ingredients and three modes of action, we observed widespread STB in nearly all of the experimental plots. There were obvious differences in overall levels of STB infection on different cultivars. Comparison of overall levels of STB disease with nearby untreated plots showed that the fungicides significantly suppressed STB development. STB was the dominating disease in the fungicide-treated plots; other leaf diseases were present at very low levels or entirely absent as a result of the fungicide treatments. Hence, this experiment provided an unusual opportunity to assess quantitative STB resistance to infection by a natural, genetically diverse population of *Z. tritici* under conducive field conditions and in the absence of competing wheat diseases.

According to weather data collected from the Lindau weather station located about 200 m away from the field site, the weather in spring-summer 2016 was cool and rainy, conditions highly conducive to development of STB (see Fig. A2 in Appendix A.1). Average daily temperature between 1 March and 27 July was 12.5^*o*^ C and the total amount of rainfall was 1245 mm. Based on daily temperature and rainfall data, we estimate that six asexual generations of *Z. tritici* occurred during this period. Between the two leaf sampling dates *t*_1_ and *t*_2_, we estimated two asexual generations (see Appendix A.1 for details of estimation).

### An overview of the dataset

A total of 21420 leaves were included in the automated analysis pipeline, with an average of 30 leaves per plot. The total leaf area analyzed was 36.8 m^2^ of which 11.3 m^2^ was recognized as damaged by STB. The mean analyzed area of an individual leaf was 17 cm^2^. In total 2.7 million pycnidia were counted. The mean number of pycnidia within a leaf was 127. A more detailed description of the overall dataset is given in Table 2. The full dataset will be submitted to www.datadryad.org after acceptance. Correlations between the two biological replicates ranged from 0.23 to 0.66, with p-values ranging from 10^-4^ to 10^-35^ (Fig. A3, see Appendix A.3 for more details).

**Table 2:** Summary of the leaf analysis

|  | Total | Mean | Median | Maximum | Minimum |
| --- | --- | --- | --- | --- | --- |
| Measured quantities |  |  |  |  |  |
| Leaf area (mm<sup>2</sup>) | 36 874 893 | 1738 | 1701 | 3626 | 103 |
| Necrotic area (mm<sup>2</sup>) | 11 162 727 | 526 | 415 | 2519 | 0 |
| Number of pycnidia | 5 148 640 | 243 | 147 | 4828 | 0 |
| Pycnidia area (mm<sup>2</sup>) | 77 681 | 3.66 | 2.35 | 56 | 0 |
| Derived quantities |  |  |  |  |  |
| PLACL (%) |  | 32 | 24 | 99.6 | 0 |
| $\rho_{\text{lesion}}$ | | 48 | 40 | 416 | 0 |
| $\rho_{\text{leaf}}$ | | 14 | 9 | 301 | 0 |

The distributions of the raw data points corresponding to individual leaves with respect to PLACL, *ρ*_lesion_ and *ρ*_leaf_ are shown in Figs. 2 and 3. The distributions of PLACL, *ρ*_lesion_ and *ρ*_leaf_ were non-normal and had outliers. All of these distributions were continuous, consistent with previous studies that hypothesized that the majority of STB resistance in wheat is quantitative (Stewart et al., 2016).

**Figure 2:**
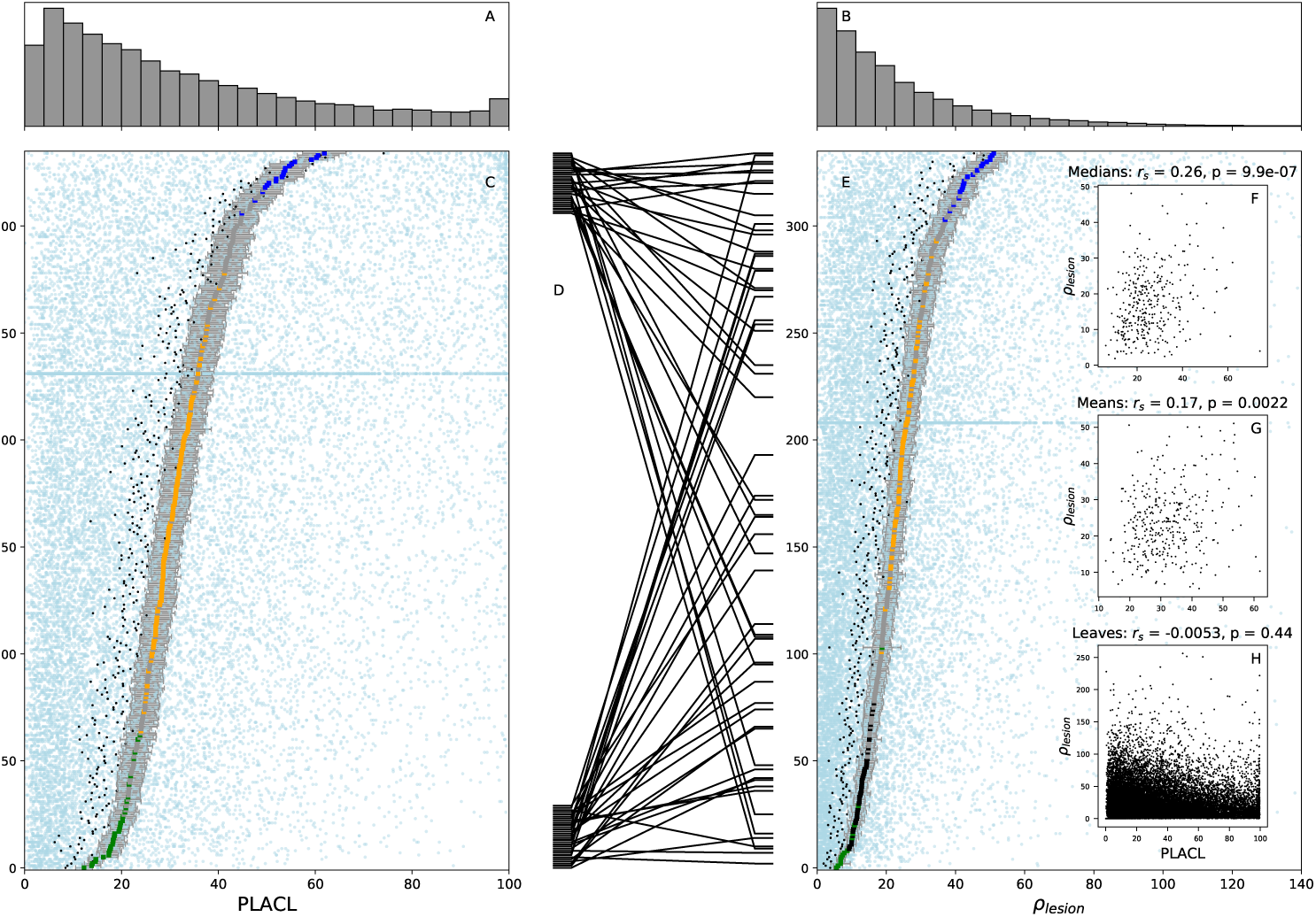
Ranking of wheat cultivars according to their resistance against host damage (panel C) and pathogen reproduction (panel E). Wheat cultivars were ranked in order of increasing susceptibility to STB based on their mean values of PLACL (percentage of leaf area covered by lesions, panel C) and *ρ*_lesion_ (pycnidia density per lesion, panel E). Colored markers depict means over two replicates and two time points, error bars indicate standard errors of means. Different colors of markers (blue, orange, black, green) represent significantly different groups of cultivars based on Kruskal-Wallis multiple comparison FDR correction, gray dots correspond to cultivars that could not be assigned to any group. Light blue dots represent average values for individual leaves. Cultivar CH Claro appears as a horizontal blue line due to its 21-fold higher number of replicates. Black dots show medians for each cultivar. Lines in panel D represent changes in ranking between PLACL and *ρ*_lesion_ among the 30 most susceptible and the 30 most resistant cultivars based on PLACL. Panels A and B show frequency histograms of PLACL and *ρ*_lesion_ based on individual leaf data from the two replicates and two time points. Three insets in panel E illustrate the relationship between PLACL and *ρ*_lesion_ based on individual leaf data (panel H), means (G) and medians (panel F). Panels B and E extend only up to 140 along the *x*-axis, missing 90 data points with values between 140 and 256.

**Figure 3:**
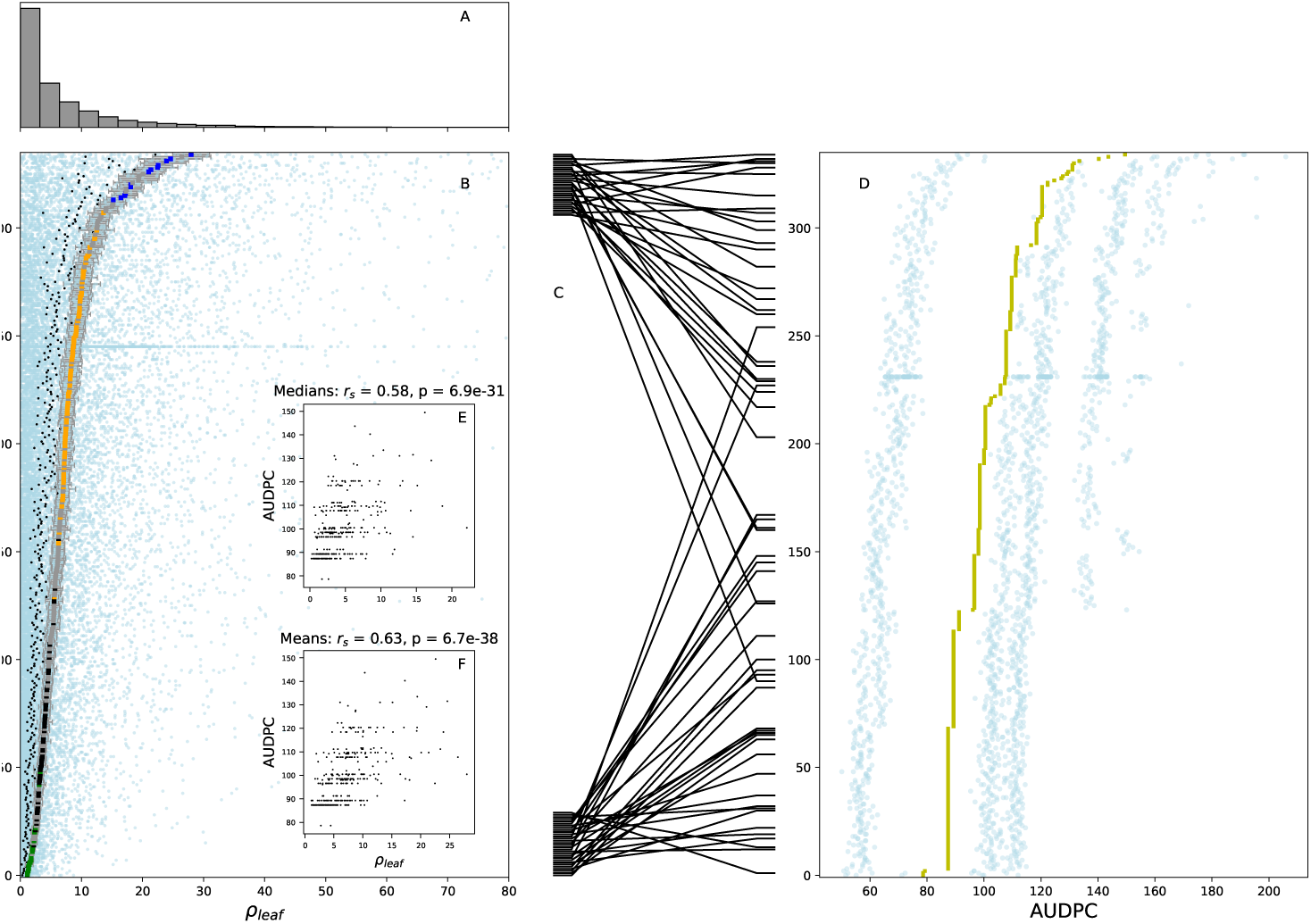
Ranking of wheat cultivars according to the resistance measure, *ρ*_leaf_, that combines host damage and pathogen reproduction (panel B) and according to AUDPC (area under the disease progress curve) based on conventional visual assessments (panel D). In panels B and D cultivars are ranked in order of increasing susceptibility based on mean values of leaf pycnidia density, *ρ*_leaf_ (panel B), and AUDPC (panel D). Colored markers show means over two replicates and two time points, error bars indicate standard errors of means. Different colors of markers (blue, orange, black, green) represent significantly different groups of cultivars based on a Kruskal-Wallis multiple comparison with FDR correction, gray dots correspond to cultivars that could not be assigned to any group. Light blue dots represent average values for individual leaves. Cultivar CH Claro appears as a horizontal blue line due to its 21-fold higher number of replicates. Black dots show medians for each cultivar. Lines in panel C represent changes in ranking between *ρ*_leaf_ and AUDPC among the 30 most susceptible and the 30 most resistant cultivars found in *ρ*_leaf_. Panel A shows a frequency histogram of *ρ*_leaf_ based on individual leaves from two replicates and two time points. Two insets illustrate the relationship between *ρ*_leaf_ and AUDPC with respect to means (panel F) and medians (panel E). Panels A and B extend only up to 80 along the *x*-axis, missing 63 data points with values between 80 and 244. Gaussian noise with variance *σ*^2^ = 4 was added to raw (blue) data points in panel D to improve their visibility.

For the visual assessments conducted across two replicates and three time points, the lowest score was 1 and the highest score was 4 (on a 1-9 scale). The lowest AUDPC value was 81, the highest value was 154 and the average AUDPC across all cultivars was 103. The Spearman correlation between visual scores of the two replicates was not significant in the first assessment but was significant in the second and third assessments. The visual assessment also found that yellow rust was present in about 1 % of plots on 20 May and in about 2 % of plots on 21 June, 2016; Septoria nodorum blotch was present in only a single plot on 21 June, 2016 (about 0.1 % of plots); Fusarium head blight was present in about 2 % of plots only on 21 June, 2016.

### Host damage vs. pathogen reproduction

From the raw data obtained via AIA, we derived three quantitative resistance measures: PLACL, *ρ*_lesion_ and *ρ*_leaf_. PLACL is defined as the necrotic leaf area divided by the total leaf area, *ρ*_lesion_ is the total number of pycnidia divided by the necrotic leaf area and *ρ*_leaf_ is the total number of pycnidia divided by the total leaf area. *ρ*_leaf_ can also be calculated from the first two factors as

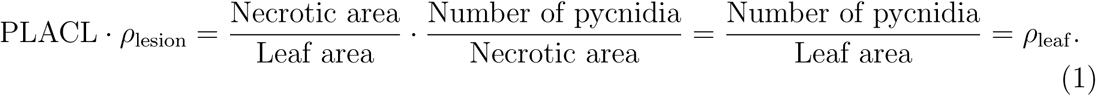

PLACL characterizes host damage due to pathogen infection while *ρ*_lesion_ characterizes pathogen reproduction on the necrotic leaf tissue. *ρ*_leaf_ is the product of these two quantities, combining host damage with pathogen reproduction. Calculation of these three quantities from the AIA data allowed us to differentiate between host damage and pathogen reproduction, and also to combine these two factors into the most integrative measure of disease intensity, providing a comprehensive insight into different components of STB resistance. Next, we ranked the wheat cultivars with respect to each of these three quantities.

### Ranking of cultivars

Resistance ranking of the cultivars was based on the three measures obtained from AIA (PLACL, *ρ*_lesion_ and *ρ*_leaf_) and the AUDPC calculated from visual scoring. For PLACL, *ρ*_lesion_ and *ρ*_leaf_, the distributions differed significantly between cultivars. For each of these three measures, the null hypothesis of identical distributions for all cultivars was rejected by a Kruskal-Wallis global comparison with *p <* 2.2 · 10^-16^. However, the global Kruskal-Wallis test did not reveal differences between distributions of the weighted visual scores (*p* = 1). Kruskal-Wallis multiple pairwise comparisons identified three significantly different groups of cultivars for PLACL and four significantly different groups of cultivars for *ρ*_lesion_ and *ρ*_leaf_ (Figs. 2 and 3). Supplemental File S3 shows that cultivar ranking based on the Kruskal-Wallis test statistic (mean ranks) is highly correlated with cultivar ranking based on the mean and median values associated with each cultivar, indicating that in the majority of cases significantly different cultivars also have different means and medians.

There were notable differences between resistance rankings based on PLACL and *ρ*_lesion_ (Fig. 2D). Several of the thirty least resistant cultivars based on host damage were ranked among the most resistant cultivars based on pathogen reproduction. Similarly, some of the most resistant cultivars based on host damage were among the least resistant cultivars based on pathogen reproduction. For example cultivar Vanilnoir showed high PLACL and low *ρ*_lesion_ whereas cultivar Rubens exhibited the opposite pattern. Visual examination of leaves belonging to cultivars that exhibited the largest difference in their ranking between PLACL and *ρ*_lesion_ confirmed these patterns. There were relatively low correlations between PLACL and *ρ*_lesion_ with respect to means (*r*_s_ = 0.17, *p* = 0.0022, Fig. 2G) and medians (*r*_s_ = 0.26, *p <* 10^-6^, Fig. 2F) taken over leaves belonging to the same cultivar. The correlation between PLACL and *ρ*_lesion_ was not significant when calculated based on individual leaf data pooled together (Fig. 2H). Supplemental Tables S3–6 supporting Fig. 2 show means, standard errors of means, medians and KruskallWallis test statistics based on PLACL and *ρ*_lesion_ for all cultivars. Supplemental File S1 gives a brief description of all supplemental files and tables.

Automated measures of quantitative STB resistance correlated strongly with the traditional measurement based on AUDPC of visual scores. Medians of PLACL and *ρ*_lesion_ correlated significantly (*p <* 10^-12^) with the AUDPC (*r*_*s*_ = 0.38 and *r*_*s*_ = 0.53 respectively). Correlations between AUDPC and means were somewhat weaker but also significant. The strongest correlation was between the combined measure *ρ*_leaf_ and the AUDPC (cf. Fig. 3, means: *r*_*s*_ = 0.63, medians: *r*_*s*_ = 0.58). More figures of cultivar ranking that support Figs. 2 and 3 with different combinations of measures are shown in Supplemental File S2. Supplemental Tables 7–8 supporting Fig. 3 show means, standard errors of means, medians and Kruskall-Wallis test statistics based on *ρ*_leaf_ for all cultivars.

### Predictors of epidemic development

So far we analyzed data for each cultivar based on pooling the sampling dates *t*_1_ and *t*_2_. Next we consider data from the sampling dates *t*_1_ and *t*_2_ separately. An important question is: To what extent can we predict a measure of disease at *t*_2_ from measurements made at *t*_1_? We address this question by investigating correlations between *t*_1_ and *t*_2_ with respect to each of the three measures: PLACL, *ρ*_lesion_, and *ρ*_leaf_ (Fig. 4). A higher degree of correlation corresponds to a higher predictive power.

**Figure 4:**
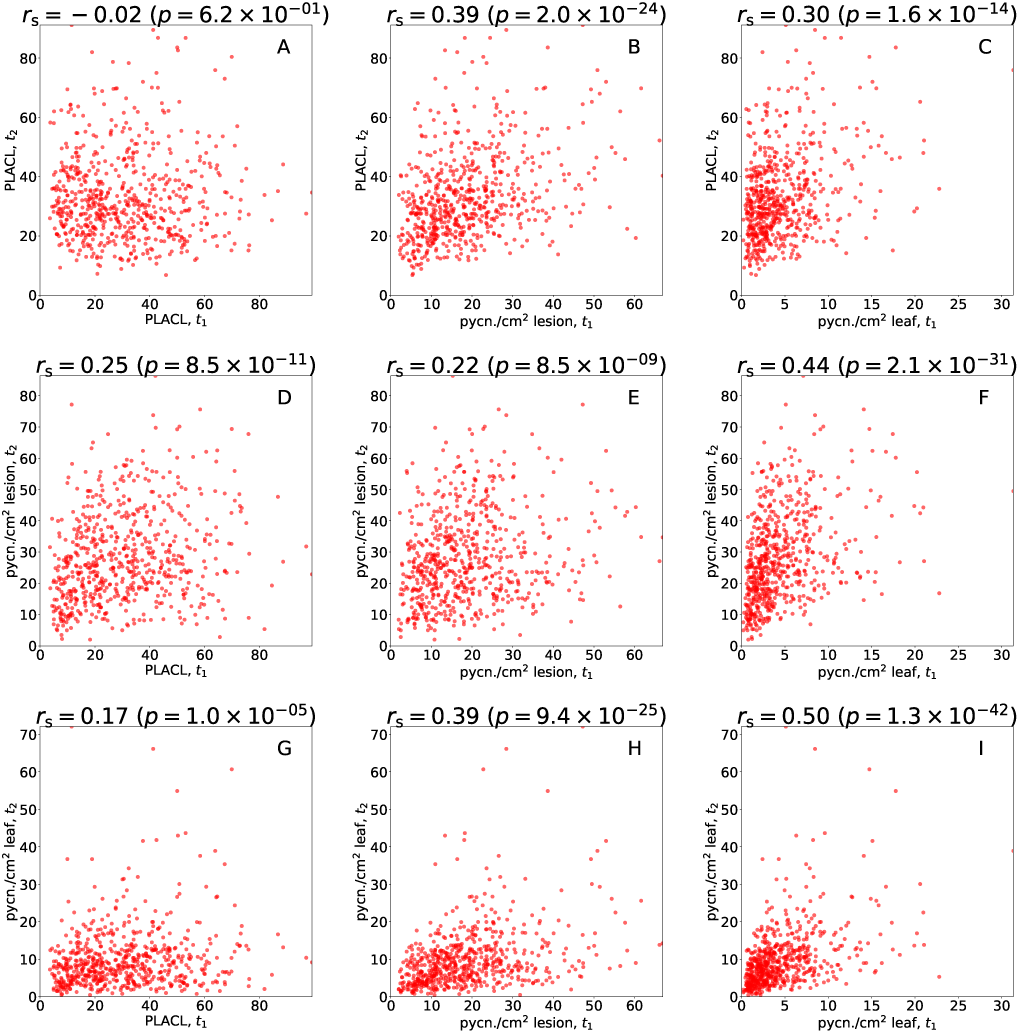
Correlations of measures of quantitative STB resistance between the first time point (*t*_1_) and the second time point (*t*_2_). The degree of correlation was quantified using Spearman’s correlation coefficient, *r*_s_. Each data point represents an average over about 15 leaves within an individual plot.

Consider the first column in Fig. 4 that illustrates how PLACL in *t*_1_ correlates with PLACL, *ρ*_lesion_ and *ρ*_leaf_ in *t*_2_. PLACL in *t*_1_ correlates somewhat better with *ρ*_lesion_ in *t*_2_ than with *ρ*_leaf_ or PLACL in *t*_2_. However, PLACL in *t*_1_ is a poorer predictor for the three quantities in *t*_2_ than the quantities that include pycnidia counts, *ρ*_lesion_ and *ρ*_leaf_ (compare the first column with the second and third columns in Fig. 4). The highest correlations emerge between *ρ*_leaf_ in *t*_1_ and *ρ*_lesion_ in *t*_2_ (*r*_s_ = 0.44) and between *ρ*_leaf_ in *t*_1_ and *ρ*_leaf_ in *t*_2_ (*r*_s_ = 0.50). As shown in the first row in Fig. 4, the best predictor for PLACL (the measure of host damage that is most likely to reflect decreased yield) in *t*_2_ is *ρ*_lesion_ (the most inclusive measure of pathogen reproduction) in *t*_1_.

Figure 4 gives a general account of the correlations/predictive power among the measured quantities in *t*_1_ and *t*_2_. We investigate more subtle patterns of this comparison in supplemental File S4, where we separate the effect of cultivar differences from the overall effect.

For a subset of 39 cultivars, we performed similar measurements in a preliminary experiment conducted in 2015 (published in Stewart et al., 2016). Comparisons between the outcomes in 2016 and 2015 indicated that PLACL exhibited no significant correlation, while *ρ*_lesion_ and *ρ*_leaf_ showed moderate and significant correlations (*r*_s_ = 0.46, *p* = 0.0035 for *ρ*_lesion_ and *r*_s_ = 0.41, *p* = 0.0092 for *ρ*_leaf_). This further highlights the importance of measuring quantitative resistance based on pathogen reproduction as it provided a more consistent measure of STB resistance between years and environments than measures of STB resistance based on host damage.

### Increase of resistance to STB between

*t*_1_ **and** *t*_2_. Next we investigate the difference between *t*_1_ and *t*_2_ to identify cultivars that exhibit an increase in resistance over time. We will refer to this as “late-onset” resistance. Each of the quantities, PLACL, *ρ*_lesion_ and *ρ*_leaf_, increased on average between the two time points (Fig. 5). The difference was somewhat larger for *ρ*_leaf_ than for PLACL and *ρ*_lesion_. The overall mean differences are smaller than the variance of differences in individual cultivars. This is because positive changes in some cultivars were compensated by negative changes in other cultivars (as seen in Fig. 5).

**Figure 5:**
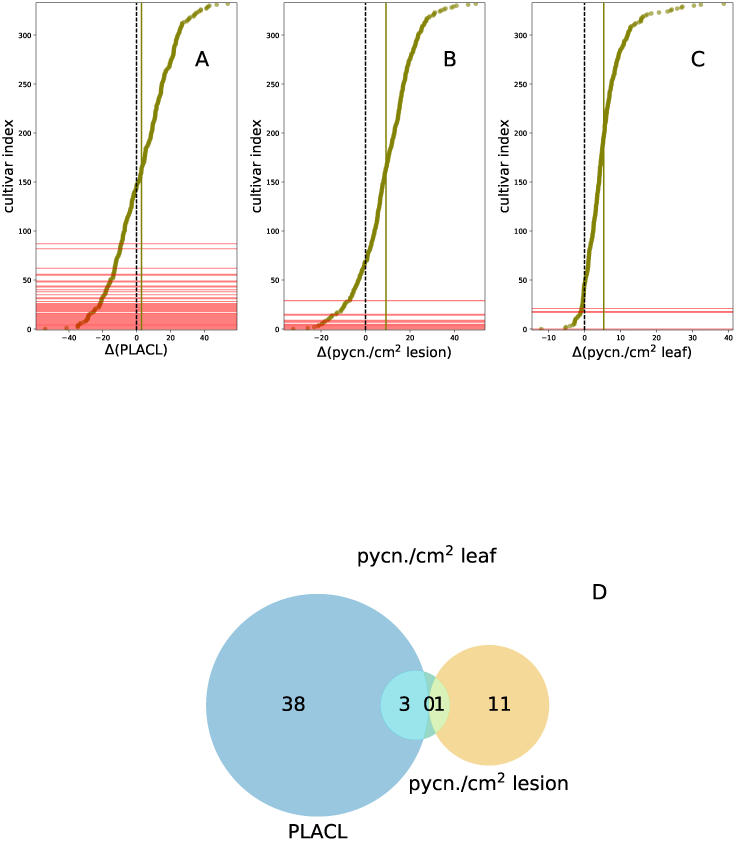
Indicators of late-onset STB resistance. *x*-axes show differences between mean values at time point *t*_2_ and time point *t*_1_ for the three quantities (yellow markers): PLACL (panel A), *ρ*_lesion_ (panel B) and *ρ*_leaf_ (panel C). *y*-axes represent cultivar indices; cultivars are sorted in order of increasing differences. Solid vertical lines show differences averaged over all cultivars; dashed vertical lines show zero differences. Cultivars exhibiting significant negative differences (according to Wilcoxon rank sum test with the FDR correction, *p <* 0.01) are shown in red. Panel D depicts a Venn diagram of cultivars with significant negative differences with respect to the three measures of resistance.

We investigated the negative changes in more detail. We identified 53 cultivars in total (marked in red in Fig. 5) that exhibited significant negative changes with respect to at least one of the quantities: PLACL, *ρ*_lesion_ or *ρ*_leaf_. We see from the Venn diagram in Fig. 5D that the number of cultivars showing significant negative change in PLACL (41 cultivars) is higher than in *ρ*_lesion_ (12 cultivars). Four cultivars showed significant negative change with respect to *ρ*_leaf_. The number of cultivars exhibiting significant negative change in terms of only one quantity were: 38 for PLACL, 11 for *ρ*_lesion_ and zero for *ρ*_leaf_. Interestingly, none of the cultivars exhibited significant negative change for both PLACL and *ρ*_lesion_. PLACL decreased the most in cultivars Achat and Mewa (by 54 and 41 units, respectively). *ρ*_lesion_ decreased the most in cultivars Parador and Cetus (by about 33 and 26 units, respectively). *ρ*_leaf_ decreased the most in cultivars Cetus and Urban (by 12 units and 1 unit, respectively). Lists of cultivars with their corresponding magnitudes of changes are given in Supplemental Tables S9–11.

## Discussion

### Novel aspects of the experimental design

Although fungicides suppressed STB development as compared to untreated plots, the most important benefit of the fungicide applications in the context of this experiment was to eliminate competing diseases (rusts, powdery mildew, tan spot, septoria nodorum blotch) that usually co-exist with STB in naturally infected fields. This resulted in a nearly pure culture of STB across both replications. Virtually every disease lesion found on a leaf was shown to be the result of an infection by *Z. tritici*. The widespread STB infection found in this experiment could be explained by the cool and wet weather during the 2015-2016 growing season that was highly conducive to development of STB coupled with a significant amount of resistance to azoles in European populations of *Z. tritici* (Brunner et al., 2008).

This experimental design provided an unusual opportunity to directly compare levels and development of STB infection and STB resistance across a broad cross-section of elite European winter wheat cultivars. Combined with the novel automated image analysis method, this allowed us to collect a large amount of high-quality data with a relatively low workload.

The measures of resistance that we characterized here, on average, represent “general” or “field” resistance because the experiment was conducted using natural infection in a year that was highly conducive to STB. Cultivars that were highly resistant under these conditions are more likely to be broadly resistant when exposed to the typical genetically diverse *Z. tritici* populations (Linde et al., 2002; Zhan et al., 2003).

### Comparison between datasets obtained from automated image analysis and visual scoring

Conventional visual assessment typically quantifies leaf necrosis (host damage) caused by the pathogen by integrating disease severity (as PLACL) and incidence (as the proportion of leaves having necrotic lesions) into a single index. Pycnidia (an indicator of pathogen reproduction) are typically considered as a presence/absence qualitative trait that helps to separate STB lesions from lesions caused by other leaf spotting diseases (e.g. tan spot or septoria nodorum blotch) occurring on the same plants. The conventional visual assessment is fast: only about nine hours in total were needed to assess more than 700 small plots three times during the season by a single person (i. e., about three hours at each measurement date).

Visual scoring benefits from a large sample size, as almost all leaves in a plot are considered during a typical visual assessment compared to only 15 leaves used for image analysis. However, due to the subjective nature of the conventional scoring process, the sample size used is not clearly defined and we could not obtain statistically significant differences between cultivars based on the conventional visual scores. Moreover, assessment of changes in trait values between two time points is very difficult with conventional visual scores. Uncertainty in detection of pycnidia in the conventional measurement may lead to misclassification of resistance. The degree of STB resistance may be overestimated on cultivars that are susceptible to host damage (have high PLACL) but suppress pathogen reproduction (have low *ρ*_lesion_), because failure to detect pycnidia may be interpreted as an absence of the disease. On the other hand, if the visual assessment finds pycnidia (typically scored as present or absent), the resistance of a cultivar may be underestimated if the degree of suppression of pycnidia production is not considered.

In contrast, automated analysis of individual leaves enables independent measurement of different aspects of conditional disease intensity. The advantage of the automated method based on leaf images is that it accounts for both host damage and pathogen reproduction in a reproducible, quantitative way with a well-defined sample size. This alleviates the uncertainty in detecting pycnidia and also allowed us to find statistically significant differences between cultivars. However, this method is more labor-intensive than visual scoring. About 360 person-hours were needed to collect and process *≈*22000 leaves and obtain the raw data.

Although this is an automated method, one needs to carefully determine its error rate at every use (as we described in the Materials and Methods), because errors may be cultivar-specific and also depend on environmental conditions. Sources of errors can be minimized in the future by further improving the experimental methodology. For example, leaves with severe defects can be excluded at the stage of mounting on paper sheets, scanning errors can be minimized by introducing an additional control step after scanning, and collector bias can be minimized by improved training of leaf collectors. Importantly, we plan to further minimize current deficiencies in pycnidia and lesion detection by incorporating the image analysis into a machine learning algorithm.

Manual generation of this large dataset that included more than two million pycnidia would not be practical. However, we demonstrated that combining cheap flatbed scanners with public domain software to conduct automated image analysis makes it feasible to separate different components of epidemic outcome affected by presumably different components of quantitative host resistance on a large scale.

### Host damage vs. pathogen reproduction

Biologically, PLACL reflects the pathogen’s ability to invade and damage (necrotize) the host leaf tissue, while *ρ*_lesion_ reflects the pathogen’s ability to convert the damaged host tissue into reproductive structures and eventually into offspring. From the host’s perspective, PLACL can be interpreted as the degree of susceptibility to damage caused by the pathogen during the infection process (e. g., through secretion of phytotoxins) or by host defense reactions (e. g., the hypersensitive response) activated after detecting the pathogen. *ρ*_lesion_ can be interpreted as measuring the host’s ability to suppress pathogen reproduction: a more susceptible host enables more pathogen reproduction per unit of infected leaf area. Automated image analysis enabled us to differentiate between host damage and pathogen reproduction and hence to measure these as separate components of as STB epidemic that reflects differences in components of resistance between wheat cultivars.

The phenotypic differences observed in our experiment may reflect different sets of genes underlying the two resistance traits. We hypothesize that PLACL reflects the additive actions of toxin sensitivity genes carried by different wheat cultivars that interact with host-specific toxins produced by the pathogen, as shown for *Parastagonospora nodorum* on wheat (e. g. Friesen et al., 2008; Oliver et al., 2012). We hypothesize that pycnidia density reflects additive actions of quantitative resistance genes that recognize pathogen effectors (e.g. *AvrStb6* recognized by *Stb6*; Zhong et al. (2017)) and activate plant defenses in a quantitative way (Krattinger et al., 2009). Another possibility is that pycnidia density reflects the availability of nutrients or the concentrations of antimicrobial compounds present in the necrotic host tissue. We anticipate testing these hypotheses by combining the phenotypic data reported here with wheat genome data (International Wheat Genome Sequencing Consortium, 2014) to conduct a genome-wide association study (GWAS) aiming to identify chromosomal regions and candidate genes underlying these components of quantitative STB resistance that may explain the observed differences in epidemics.

The density of pycnidia per unit leaf area, *ρ*_leaf_, is a measure of disease intensity that incorporates both host damage and pathogen reproduction, reflecting the pathogen’s overall ability to convert healthy host tissue into reproductive units that can drive a new cycle of infection. The complex nature of quantitative host-pathogen interactions may lead to high PLACL combined with low *ρ*_lesion_ or low PLACL combined with high *ρ*_lesion_. In both of these cases, the infection is less severe than when an interaction leads to a high *ρ*_leaf_. Therefore, we believe that *ρ*_leaf_ better characterizes overall host resistance, pathogen fitness and STB intensity than PLACL or *ρ*_lesion_.

### Ranking of cultivars

The overall differences among cultivars were relatively small, except for a small number of cultivars at each extreme. These small differences are reflected in the limited number of significantly different groups of cultivars. However, our analyses showed that ranking of cultivars differs considerably with respect to the two measures of quantitative resistance: resistance to host damage and resistance to pathogen reproduction (Fig. 2D). More specifically, we identified cultivars that strongly exhibit one component of epidemic outcome, while the other component is virtually absent (points in the upper-left and lower-right corners of Fig. 2F). We expect that resistance that lowers pycnidia production will have a greater overall impact on reducing damaging STB epidemics because it will reduce the rate of epidemic increase more than resistance to host damage. Conventional phenotyping based on visual assessment does not enable separation of these different components of epidemic outcome and the underlying components of resistance.

We expected that conventional visual assessment (based on AUDPC) would correlate best with the measurement of host damage (PLACL), as conventional assessment consists mainly of quantifying leaf necrosis, while using pycnidia mainly to confirm the presence of STB. Surprisingly, the AIA measure that combines host damage and pathogen reproduction (*ρ*_leaf_) gave the best correlation with the conventional visual assessment. A possible explanation for this high correlation is that the conventional assessment may actually quantify pycnidia, but in a subjective way. An alternative explanation is that because conventional assessment includes both disease severity and incidence, it captures the overall pathogen population size that depends on both host damage and pathogen reproduction.

The correlation between our combined measure (*ρ*_leaf_) and the conventional measure (AUDPC) in Fig. 3 indicates that breeders may have selected for cultivars that combine resistance to host damage and resistance to pathogen reproduction. This hypothesis is supported by the small but significant positive correlations between PLACL and *ρ*_lesion_ in terms of both means and medians over individual cultivars (Fig. 2F). See Appendix A.4 for a discussion on the use of means vs. medians in the analysis.

Our rankings are based on conditional measures of disease intensity and do not take into account resistance that might lead to a decreased number of diseased leaves. In order to include this aspect of resistance in future rankings, our conditional measurements will need to be complemented by measurements of STB incidence.

### Predictors of epidemic development

PLACL on the upper leaves is a key determinant of disease-induced yield loss (e.g., Brokenshire, 1976). Hence, the ability to predict PLACL on the upper leaves late in the growing season based on an early season measurement may improve disease control. Our results suggest this possibility: the measure that quantifies pathogen reproduction, *ρ*_lesion_, evaluated early in the season (at *t*_1_, approximately GS 41) predicts the late-season host damage (PLACL at *t*_2_, approximately GS 75-85) better than early-season PLACL or *ρ*_leaf_ (see Fig. 4, compare panel B with panels A and C). We postulate that this finding could improve decision-making for fungicide application: one may need to apply fungicides only if *ρ*_lesion_ exceeds a certain threshold early in the season.

### Increase of resistance to STB between *t*_1_ and *t*_2_

Negative changes with respect to traits characterizing host susceptibility (PLACL, *ρ*_lesion_, and *ρ*_leaf_) suggest an increase in host resistance over time. We found that none of the cultivars exhibited this property for both PLACL and *ρ*_lesion_. Accordingly, none of the cultivars showed a significant decrease only for the combined measure *ρ*_leaf_. This may indicate that the genetic basis of the late-onset resistance differs for host damage and pathogen reproduction. These outcomes may help to reveal the genetic basis of “late-onset” resistance to STB (e.g. using GWAS or QTL mapping).

### Conclusions

We utilized a novel phenotyping technology based on automated analysis of digital leaf images to compare quantitative resistance to STB in 335 European wheat cultivars naturally infected by a highly variable local population of *Z. tritici*. This method allowed us to distinguish between resistance components affecting host damage (PLACL) and resistance components affecting pathogen reproduction (*ρ*_lesion_). Since measurements of pycnidia density cannot be accomplished on such a large scale with traditional assessment methods, our new method provides a powerful tool for measuring quantitative resistance to STB.

As suggested by Simko et al. (2017), digital phenotyping reduces subjectivity in trait quantification and may reveal small but meaningful differences that would not be observed with conventional visual assessment. Further development of this method could involve adjustment of the analysis parameters to be optimized for each cultivar separately, machine learning for more precise detection of symptoms and combining it with incidence data gathered by image analysis of high-quality canopy images produced by devices like the phenomobile (Deery et al., 2014) or the ETH field phenotyping platform (Kirchgessner et al., 2017).

### Outlook

Our approach can be readily applied to classical phenotype-based selection and breeding. Cultivars that show high resistance to both host damage and pathogen reproduction will be most likely to strongly suppress the pathogen population at the field level and result in less overall damage due to STB. Importantly, cultivars that show the highest resistance based on either host damage or pathogen reproduction can be used in breeding programs as independent sources of different components of resistance. We believe our approach provides a powerful method to specifically breed wheat cultivars carrying resistance that suppresses pathogen reproduction.

While the ability to separate phenotypes associated with two different aspects of resistance provides new avenues for resistance breeding, our hypothesis that the components of resistance explaining the two components of epidemic outcome are under separate genetic control remains to be confirmed by further research. We anticipate that future genetic studies (e.g. using GWAS) based on these phenotypic data will enable us to identify genetic markers that are linked to the different types of resistance. These markers could then enable joint selection of the different forms of resistance via marker-assisted breeding or in a genomic selection pipeline. If we can validate our hypothesis that toxin sensitivity genes underlie differences in PLACL among cultivars, the breeding objective would be to remove these sensitivity genes (Friesen et al., 2008; Oliver et al., 2012).

## Acknowledgements

PK and AM gratefully acknowledge financial support from the Swiss National Science Foundation through the Ambizione grant PZOOP3 161453. The authors are grateful to Danilo Dos Santos Pereira, Simone Fouche and Lukas Meile for help in collecting and processing leaf samples, to Ethan Stewart for advice and guidance in using the leaf image analysis software, and to Hansueli Zellweger for managing the wheat trial.

## A. Appendix

### A.1. Estimation of the number of pathogen generations

We estimated the number of asexual cycles of pathogen reproduction (number of generations) using data on the dependency of the latent period of *Z. tritici* on temperature Fig. 5 in Shaw (1990). We are interested in the overall relationship between the latent period and the temperature and would like to use the largest amount of data available. For this reason, we pooled together the data available for two cultivars, Avalon and Longbow, recorded by Shaw (1990). Next, we fitted the polynomial function

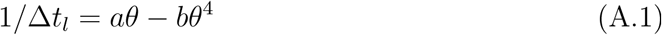

to the resulting data. Here, Δ*t*_*l*_ is the latent period, *θ* is the temperature, *a* and *b* are fitting parameters. The outcome is shown in Fig. A1. Best-fit values of parameters are:

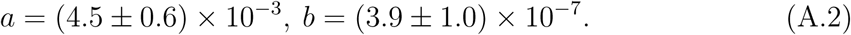

Uncertainties in Eq. A.2 represent the 95 % confidence intervals calculated from standard errors. Goodness of fit: *R*^2^ = 0.7; standard error of regression *s* = 5.5×10^-3^.

We then used average daily temperatures and the amount of rainfall measured at the Lindau weather station located close to our experimental site to estimate the number of pathogen generations, *n*_*g*_. We performed estimation of *n*_*g*_ from the period of most active vertical growth until harvest (March 1 until July 27) and between the two sampling dates (from May 1 until July 4). First, we determined the average latent period from the daily temperature averaged over the growing season using Eq. A.1 with the parameter values from Eq. A.2. This resulted in the value Δ*t*_*l*_ = 21 days. After that, we introduced a constraint on the number of pathogen generations using the rainfall data. According to our current understanding, rainfall is the most efficient way to release and disperse the asexual pycnidiospores. For this reason, we assumed that a cycle of asexual reproduction could only be completed after a day with at least 5 mm rainfall (similar to Zhan et al. (2002)). In this way we estimated an average of about six cycles of asexual reproduction between March 1 and July 27 and about two cycles between the two sampling dates *t*_1_ and *t*_2_ (Fig. A2).

**Figure A1:**
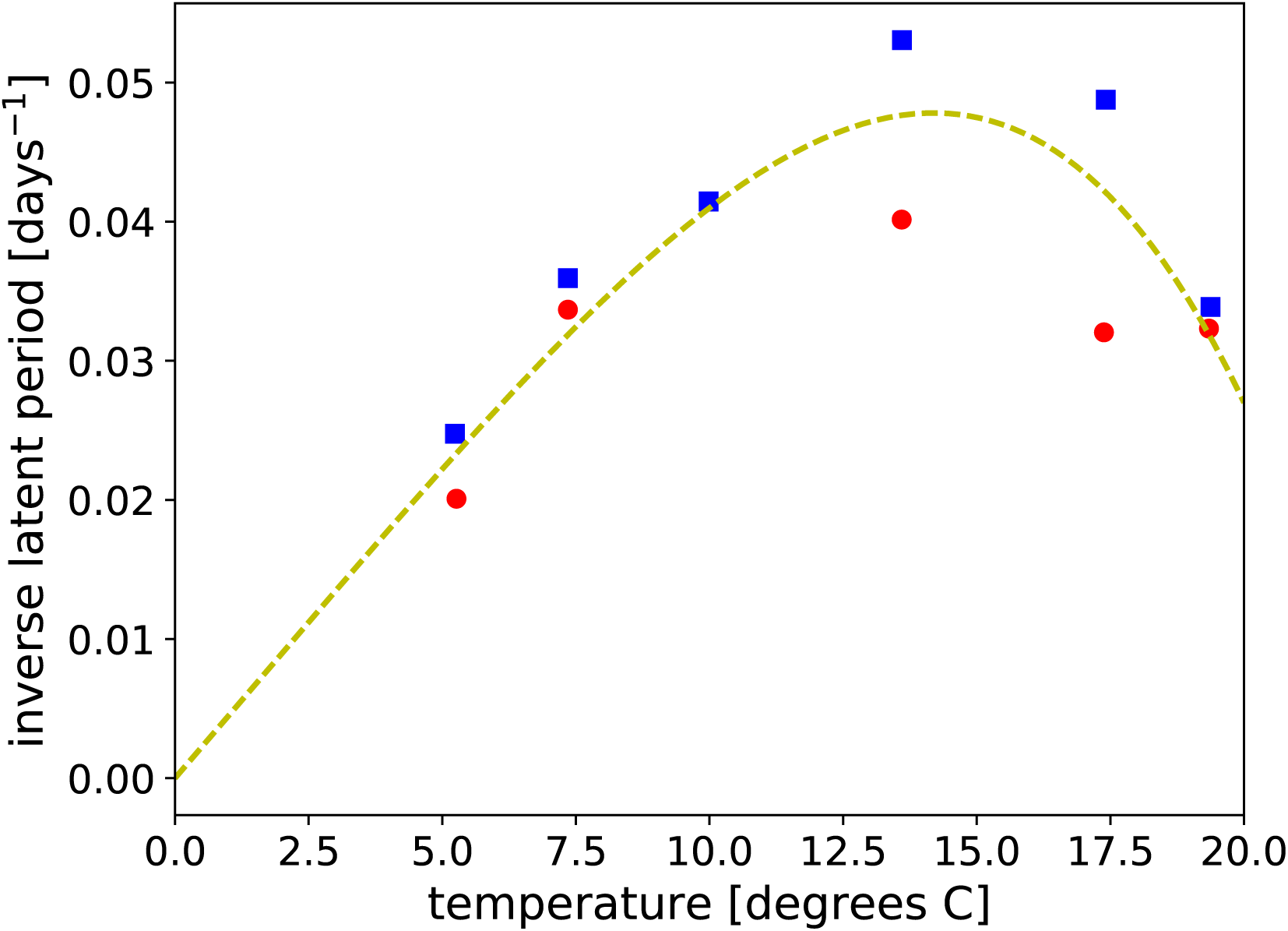
Inverse of latent period of *Z. tritici* as a function of the temperature based on data from Fig. 5 by Shaw (1990). Data from controlled-environment experiments for cultivars Longbow (circles) and Avalon (squares). Dashed curve is the best fit using the function in Eq. A.1 (see text for details).

**Figure A2:**
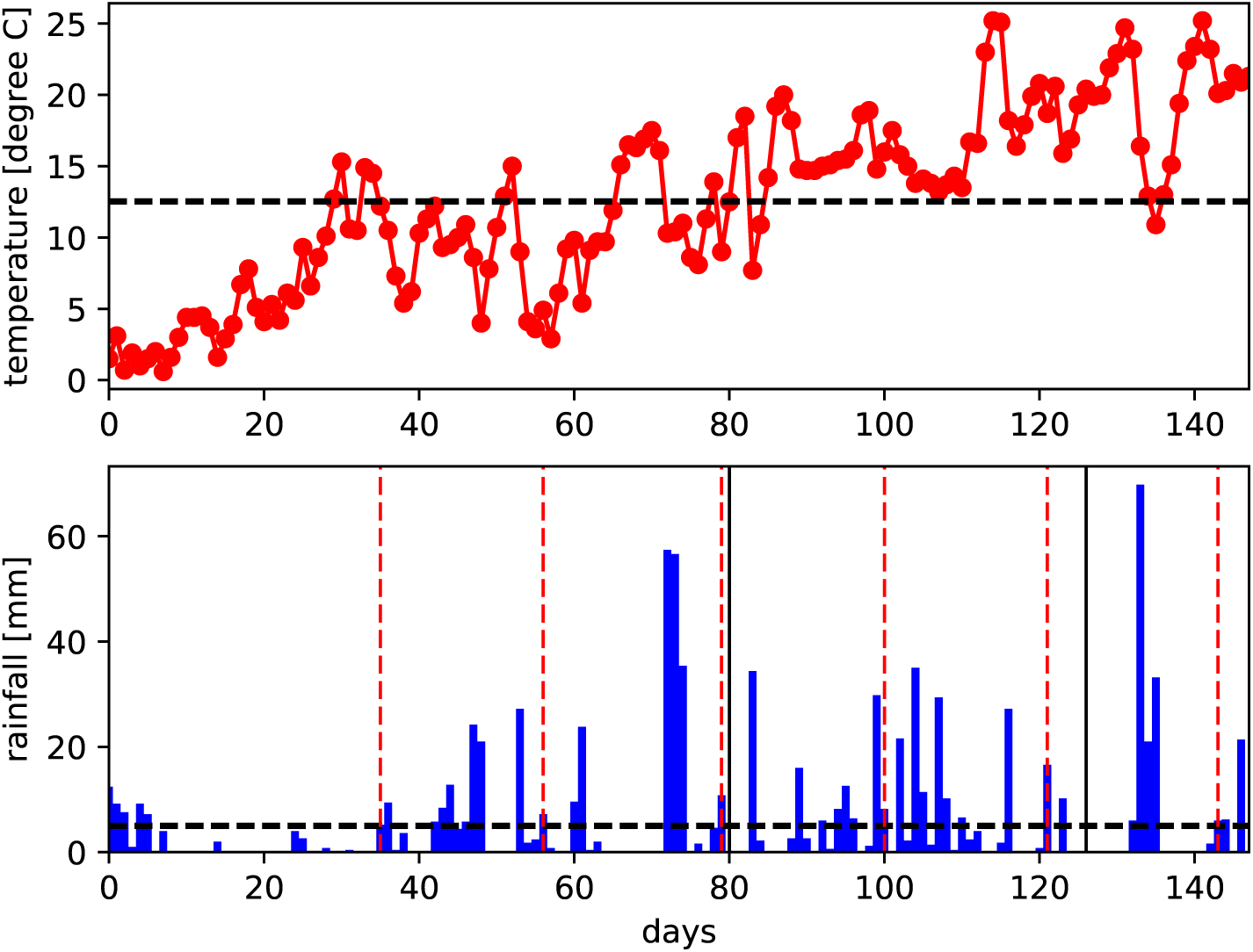
Temperature and rainfall data recorded at the Lindau weather station (data from http://www.agrometeo.ch/de/meteorology/datas) for the period between March 1 and July 27, 2016. Red vertical lines indicate the estimated generation times and black vertical lines show the sampling dates *t*_1_ and *t*_2_.

### A.2. Calculation of AUDPC based on visual assessments

We denote the values of visual scores recorded on the first, *τ*_1_, second, *τ*_2_ and third, *τ*_3_, dates (20th of May, 21th of June and 29th of June, 2016) of visual assessments as *A*_1_, *A*_2_ and *A*_3_. Area under the disease progress curve (AUDPC) was calculated for each cultivar using the visual scores in the following manner:

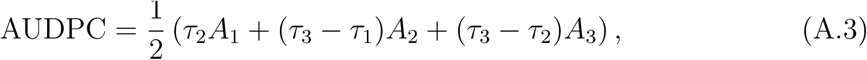

where *τ*_1_ = 14 days, *τ*_2_ = 44 days, *τ*_3_ = 52 days. Here, we assumed that disease started from zero 14 days before the first scoring, values of *A*_1_, *A*_2_ and *A*_3_ are taken as average scores over the two replicate plots. Eq. A.3 uses a trapezoidal function to interpolate between the time points in order to calculate the area under the curve. This assumes that score values are connected by linear segments. Values of AUPDC are shown for each cultivar in Fig. 3D of the main text.

To analyze differences between cultivars, we weighted scores *A*_1_, *A*_2_ and *A*_3_ in the following manner: *a*_1_ = 3*τ*_2_*A*_1_*/*2, *a*_2_ = 3(*τ*_3_ *- τ*_1_)*A*_2_*/*2, *a*_3_ = 3(*τ*_3_ *- τ*_2_)*A*_3_*/*2. The coefficients were chosen such that weighted scores *a*_1_, *a*_2_ and *a*_3_ give proportionate contribution to the AUDPC and the arithmetic average over them gives the AUDPC. The weighted scores from each replicate and each time point were given as a set of points corresponding to each cultivar (total of six measurement points per cultivar). One score was missing for one of the replicates in several cultivars. For these cultivars, we used only five measurement points for statistical analysis. We also calculated grand means over raw (not weighted) visual scores for each cultivar by taking arithmetic means over measurements in two replicates and three time points (six measurement points). Similar to the case of weighted scores, when scores were missing in one of the replicates at one of the time points, we calculated arithmetic means over the five values that were present.

Statistical differences between cultivars based on visual scoring could, in principle, be tested in a similar manner as based on image analysis data. We tested differences between distributions based on the weighted visual scores of the cultivars. The global Kruskal-Wallis test (R Core Team, 2016) revealed no differences (Kruskal-Wallis chisquared = 194.39, df = 335, *p* = 1). Surprisingly, when using unweighted visual scores there were differences between cultivars (Kruskal-Wallis chi-squared = 642.4, df = 335, *p <* 2.2 · 10^-16^). Pairwise multiple comparison with a false discovery rate p-value correction (Benjamini and Hochberg, 1995) found that the 23 most susceptible cultivars were different from the most resistant cultivar and the 155 most resistant cultivars were different from the most susceptible cultivar. However, there were 157 cultivars that were not significantly different from either extreme. The slight but significant difference between unweighted and weighted data may arise from the short time interval between *τ*_2_ and *τ*_3_. Thus cultivars cannot be distinguished by weighted scores, as the last scoring time, likely resulting in the greatest differences between visual scorings of cultivars, had the smallest effect on AUDPC and consequently differences arising from the latest scores were suppressed. Despite differences between statistical properties of unweighted and weighted data, the ranking based on them is pretty similar: Spearman’s correlation between mean ranks of unweighted and weighted scores of cultivars is high (*r*_*s*_ = 0.97, *p <* 2.2 · 10^-16^). Also Spearman’s correlations between AUDPC, mean rank of unweighted scores and arithmetic mean of unweighted scores are high (AUDPC vs. mean ranks: *r*_*s*_ = 0.97; AUDPC vs. means: *r*_*s*_ = 0.97; mean ranks vs means *r*_*s*_ = 0.998; *p <* 2.2 · 10^-16^ for each).

### A.3. Correlation between replicates

Correlations between the biological replicates ranged from 0.23 to 0.66 for AIA measurements (Fig. A3). The highest correlation between the two replicates was found in pycnidia numbers at *t*_2_. The lowest correlation was in PLACL at *t*_1_. PLACL showed the largest difference in correlation coefficients between replicates. The exceptionally low correlation between replicates at *t*_1_ for PLACL may have arisen from making the collection at a critical point in the epidemic: if the last infection cycle had been just entering the necrotic phase (as suggested in Fig. A2), there could have been a large variation between replicates by chance due to the highly variable length of the latent period for STB infection. Correlation between visual scores was not significant in first assessment but was significant and in similar range as for AIA in the second and third assessment (*r*_*s*_ = 0.37, *p* = 6 10^12^; *r*_*s*_ = 0.18, *p* = 9 10^4^, respectively).

Spearman’s correlation describes the linear relationship between rankings based on the two replicates. Moderate but highly significant correlations between replicates for each measure imply that resistance rankings based on these measures may differ considerably between replicates. We expect this to result from the shape of the quantitative resistance distributions (Fig. 2 and 3 of the main text). For all three main measures of resistance, PLACL, *ρ*_lesion_ and *ρ*_leaf_, means of the measure are quite similar for all cultivars except for a small number of cultivars at the phenotype extremes. Thus, even a small variation in these measures between replicates may result in a large change in ranking of a cultivar for one replicate to other. This is also implied by the small statistical differences between cultivars in the middle of the distributions (Fig. 2 and 3 of the main text). The same arguments hold for low (and not significant) correlations between visual scores of the replicates.

**Figure A3:**
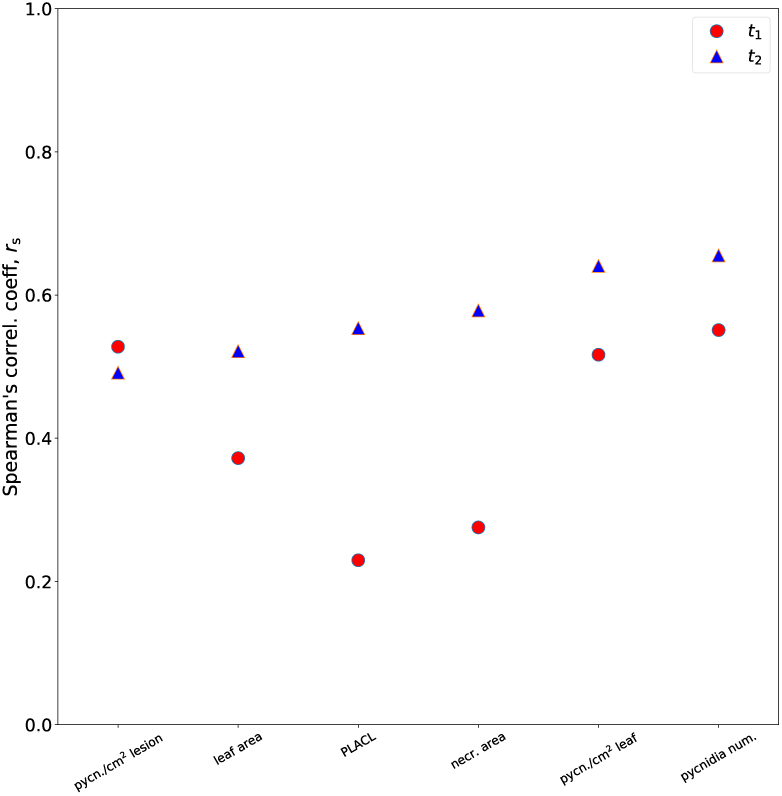
Spearman’s correlation coefficient between the mean values (taken over leaves belonging to the same cultivar) in two replicates for six different AIA measurements. All results are highly significant (*p <* 10^-7^).

### A.4. The use of means vs. medians in the analysis of STB resistance

As our data was clearly non-normal, the central tendency of the data may be better described by medians than means. Comparing medians is also biologically reasonable: total yield is more likely determined by the large number of moderately damaged wheat plants than by the few heavily diseased or dead plants. We also found stronger correlations between AUDPC and the medians of PLACL and *ρ*_lesion_ than between AUDPC and the means of PLACL and *ρ*_lesion_. Nevertheless, when comparing AUDPC and *ρ*_leaf_ we found stronger correlation between AUDPC and the means of *ρ*_leaf_ than the medians. This may indicate that mean values better describe the way breeders traditionally assess diseases caused by necrotrophs. If this insight proves to be true, it may help us to understand and counteract the subjective nature of visual assessment with its tendency to overweight fully diseased plants and thereby overestimate the overall disease severity.

## E. eXtra

### E1. Description of eXtra tables and figures

Supplemental File S1 describes content of each supplemental file and table. Supplemental File S2 shows full range versions of ranking figures (Figs. 2 and 3 of the main text) and also additional comparisons between PLACL and *ρ*_leaf_, PLACL and AUDPC, and *ρ*_lesion_ and AUDPC. Supplemental File S3 shows comparison between means, medians and mean ranks of PLACL, *ρ*_lesion_, and *ρ*_leaf_. Supplemental File S4 shows detailed analysis of predictive power of PLACL, *ρ*_lesion_, and *ρ*_leaf_ at *t*_1_ on those variables at *t*_2_ and considers separately “cultivar effect” and “pathogen effect”. Supplemental File S5 shows additional instructions and information about modifications made to the image analysis macro.

Supplemental Table S1 provides the parameter values used in the ImageJ macro and can be used directly as the input to the macro. Supplemental Table S2 defines the scale used for visual scoring of STB symptoms.

Supplemental tables supporting Fig. 2 and Fig. 3 of the main text display information on ranking of cultivars in tab-delimited text files. Supplemental Tables S3–S5 show cultivar names in first column; genetic identification number in second column; mean of observable (PLACL, *ρ*_lesion_, *ρ*_leaf_; respectively) over all the leaves of a cultivar in the third colum; standard error of the mean in the fourth column and median in the fifth column. The tables are sorted according to descending mean. In Supplemental Tables S6–S8 cultivars are ranked according to mean rank of leaves of a cultivar according to corresponding measurement. First column shows cultivar names; second genetic id; third mean rank of the data and fourth grouping code, regarding statistical differences between cultivars according to multiple pairwise Kruskal-Wallis comparison with false discovery rate p-value correction (cultivars having same letter in the grouping code are not significantly different). The tables are sorted according to descending mean rank. All tables contain a header row naming the columns.

Supplemental Tables S9–S11 supporting Fig. 5 of the main text display information on cultivars that exhibit significant negative change between *t*_1_ and *t*_2_ in mean values of PLACL, *ρ*_lesion_ and *ρ*_leaf_, respectively. Cultivars are sorted according to the difference between the mean values in *t*_1_ and *t*_2_. First column shows cultivar names; second column shows mean values in *t*_1_; third column shows mean values *t*_2_; fourth columns shows the difference between means in *t*_1_ and *t*_2_; fifth col: W statistic of the Wilcoxon rank sum test; 6th col: p-value of the Wilcoxon rank sum test with FDR correction.

### E2. Ranking of cultivars

In Fig. E2.1 cultivars are ranked as in Fig. 2 of the main text and the full range of raw data is shown for *ρ*_lesion_. Significantly differing groups of cultivars are labeled with different colors (excluding gray). Fig. E2.2 has the same organizing principle except that Fig. E2.2E shows ranking according to *ρ*_leaf_. Details of correlations are shown in insets (F, G, H) as in Fig. 2 of the main text.

In Figs. E2.3 and E2.4 the connection between traditional resistance measurement (AUDPC) and either PLACL or *ρ*_lesion_, respectively, are given similarly to what was shown in Fig. 3 of the main text. Spearman’s correlation coefficients are higher between medians of PLACL or *ρ*_lesion_ and AUDPC than between means of those and AUDPC. Details of correlations are given in the insets (E, F).

**Figure E2.1:**
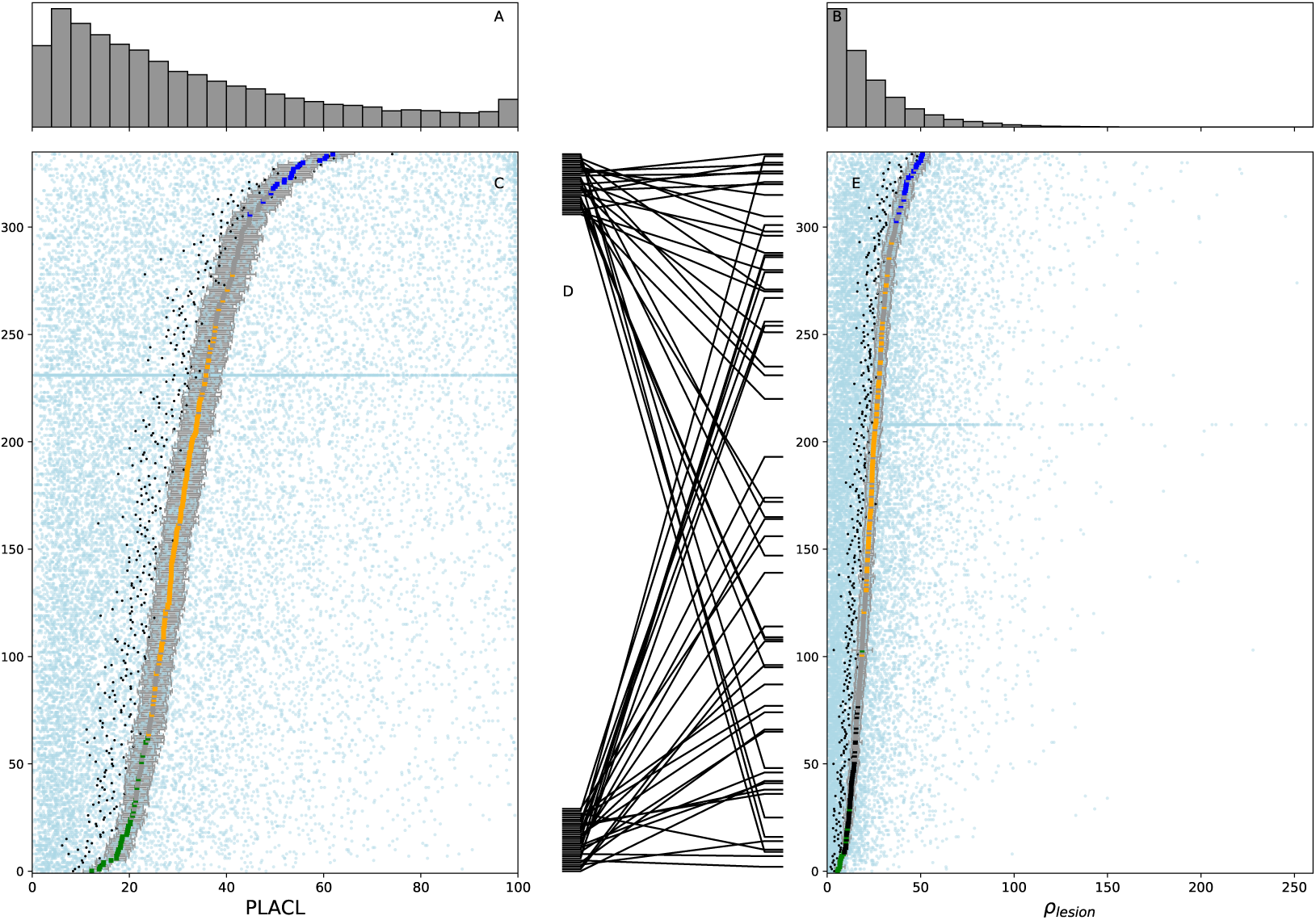
This figure is the same as Fig. 2 of the main text except that panel E shows the full range of data points of *ρ*_lesion_.

**Figure E2.2:**
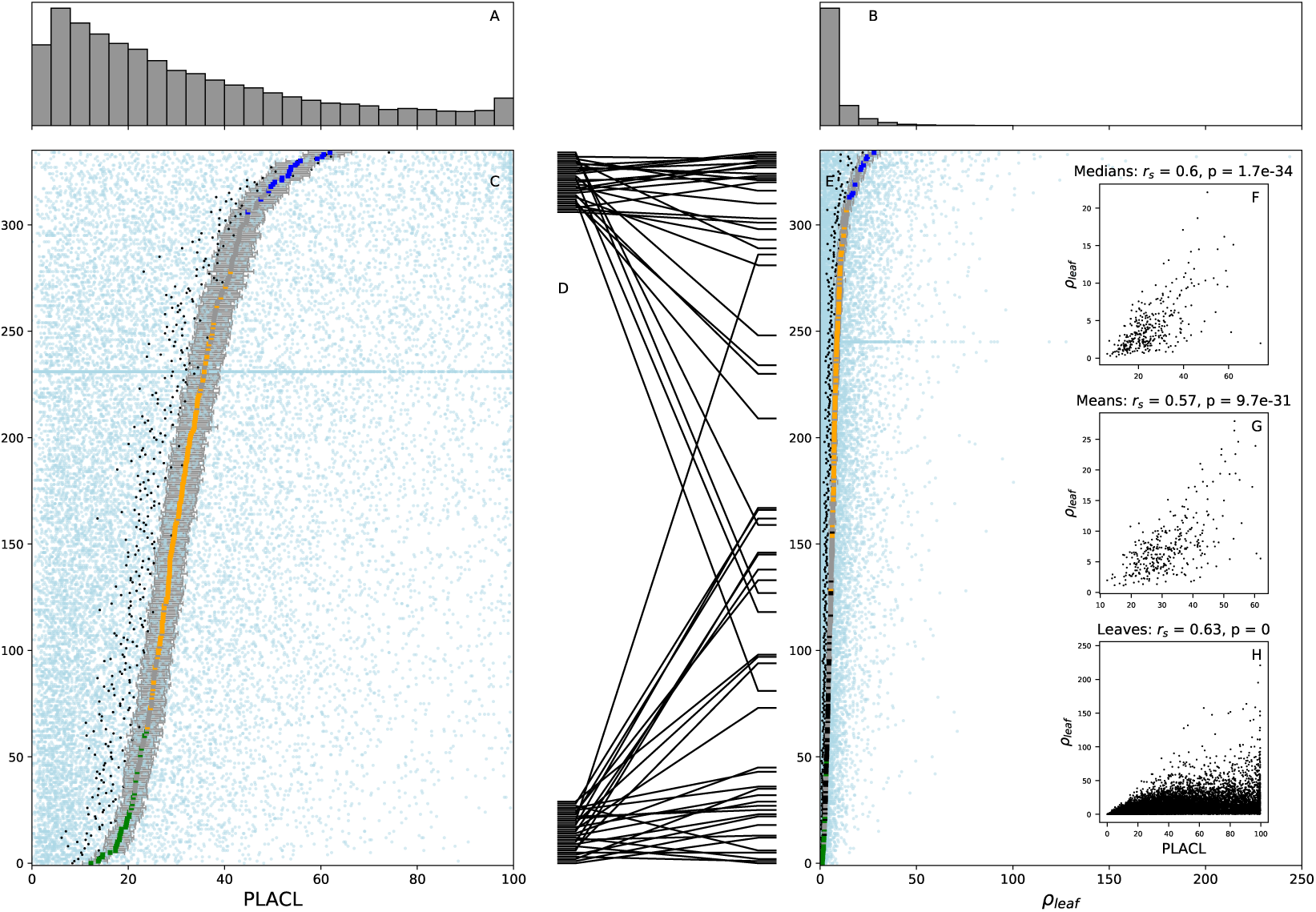
The organzation of the figure is the same as in Fig. 2 of the main text, but here panel E shows data of *ρ*_leaf_ and correspondingly insets F, G and H show correlations between PLACL and *ρ*_leaf_.

**Figure E2.3:**
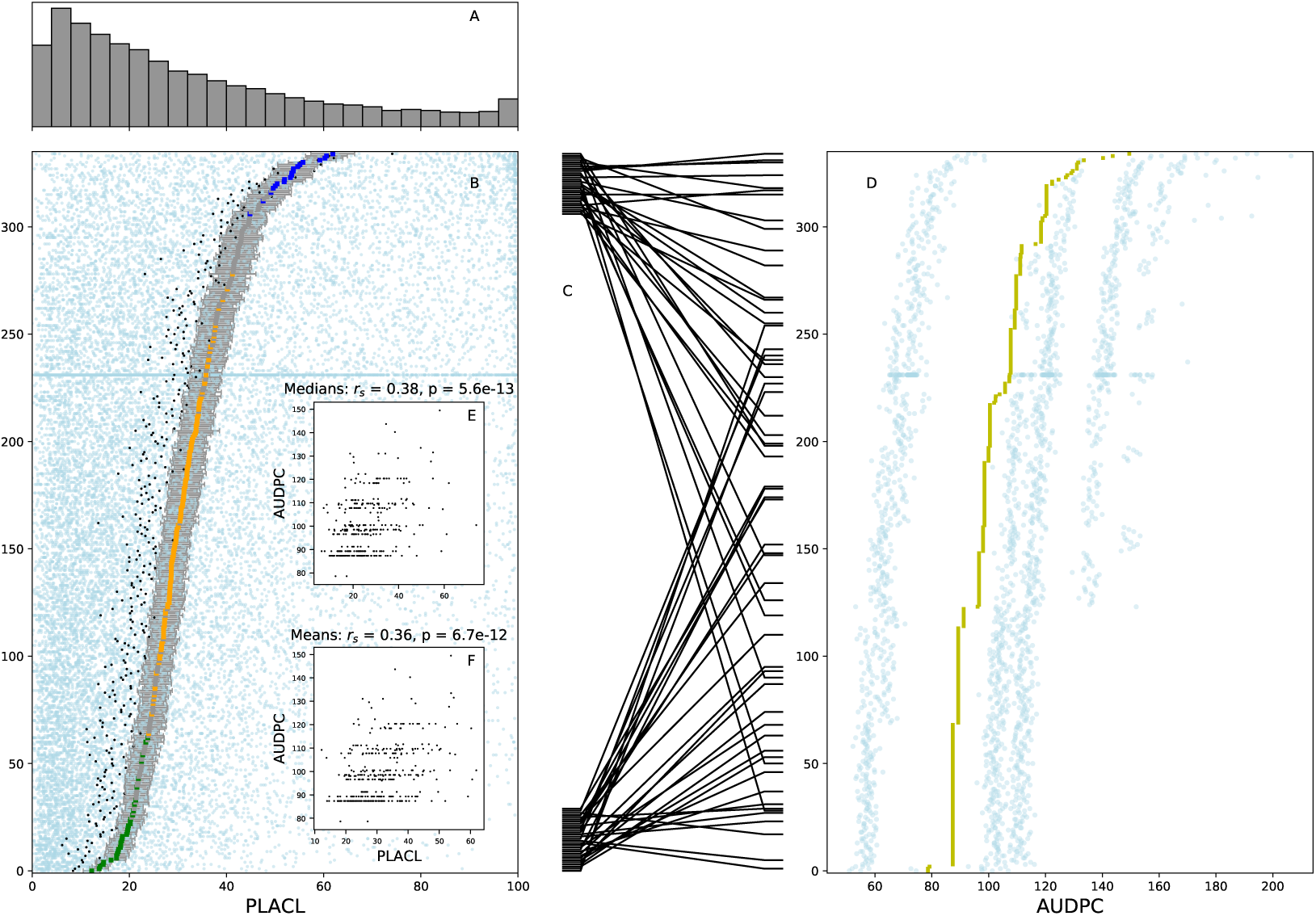
The organzation of the figure is the same as in Fig. 3 of the main text, but here panel B shows data of PLACL and correspondingly insets E and F show correlations between PLACL and AUDPC.

**Figure E2.4:**
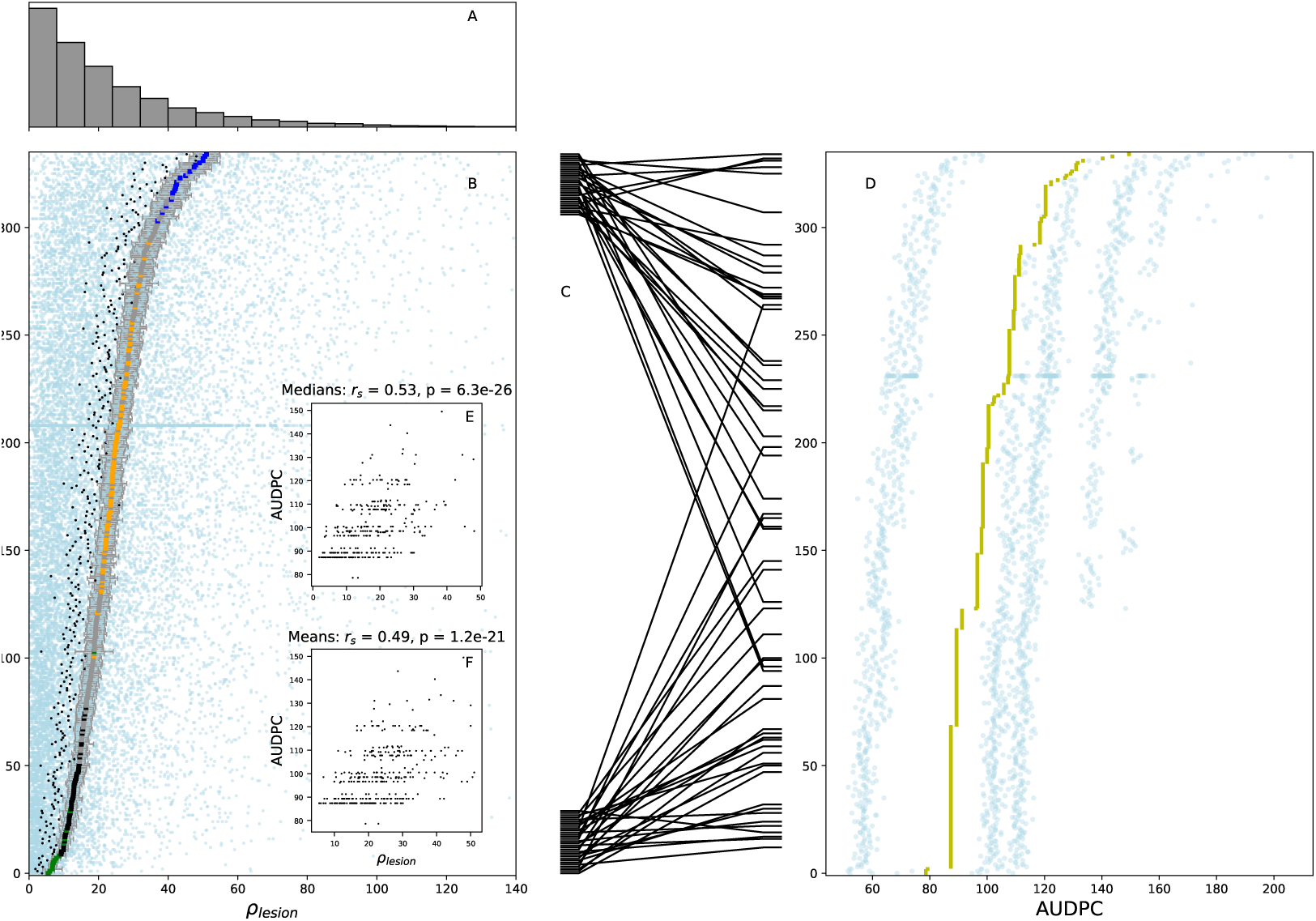
The organzation of the figure is the same as in Fig. 3 of the main text, but here panel B shows data of *ρ*_lesion_ and correspondingly insets E and F show correlations between *ρ*_lesion_ and AUDPC.

### E3. Correlation between means, medians and mean rank of PLACL, *ρ*_lesion_ **and** *ρ*_leaf_

Resistance ranking of cultivars according to means, medians and mean ranks (KruskalWallis test statistics) for PLACL, *ρ*_lesion_ and *ρ*_leaf_ are very strongly correlated (*r ≥* 0.95) (Fig. E3.1). Thus differences between mean ranks revealed by Kruskal-Wallis comparisons (cf. Fig. 2 and Fig. 3 of the main text) are correlated to differences between ranking of cultivars according to means or medians of PLACL, *ρ*_lesion_ and *ρ*_leaf_.

**Figure E3.1:**
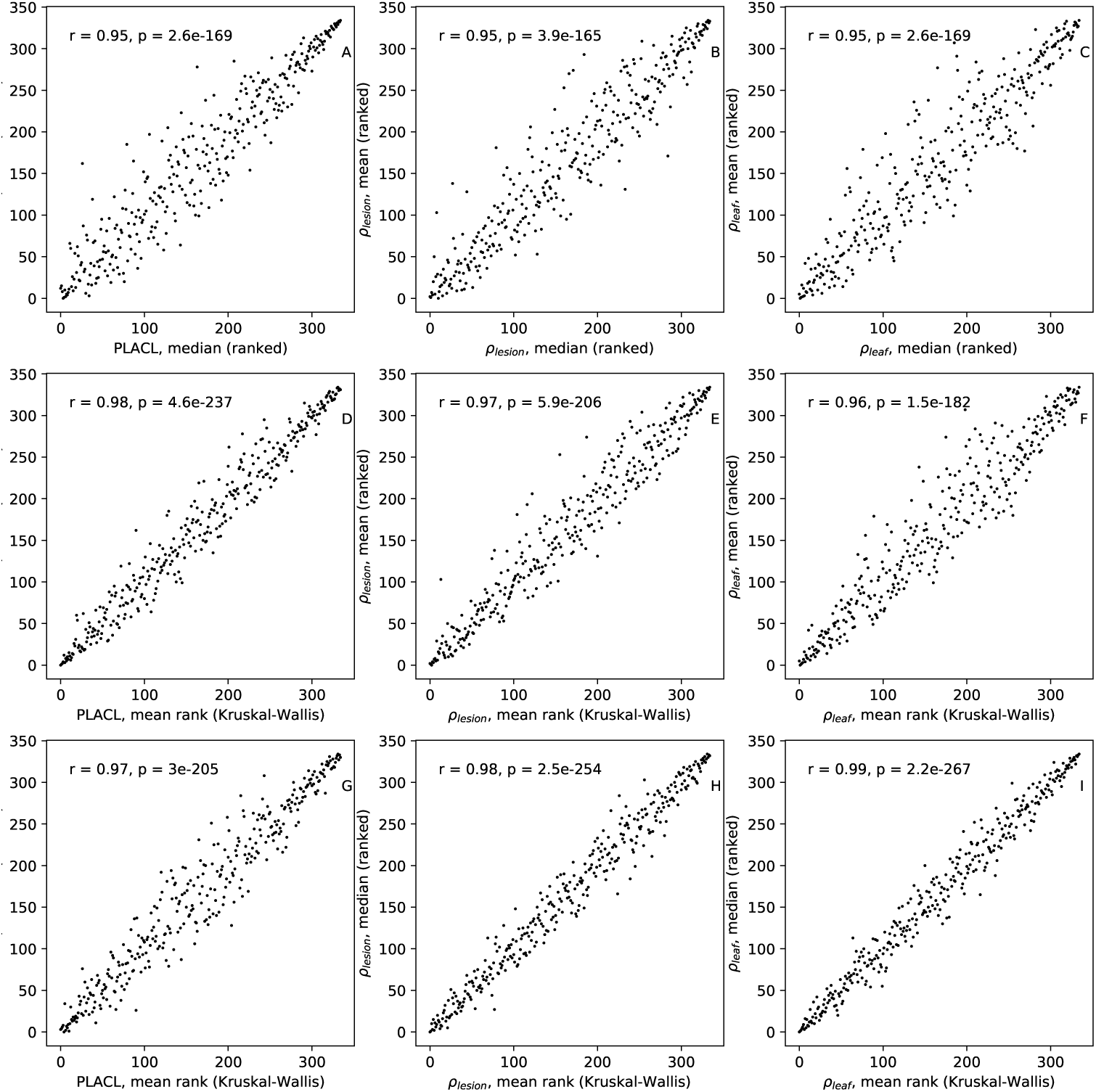
Pearson correlations between ranked means and medians and mean ranks (Kruskal-Wallis test statistics) of PLACL, *ρ*_lesion_ and *ρ*_leaf_.

### E4. Predictors of epidemic development: separating effects of cultivar and pathogen

In the section “Predictors of epidemic development” of the main text, we presented correlations between two time points for PLACL, *ρ*_lesion_ and *ρ*_leaf_ (Fig. 4 of the main text). These correlations were based on means over leaves belonging to each individual plot (with two plots per cultivar). Here, we disentangle two factors that may be responsible for the observed correlations. First is the effect of the host. In this case, the correlation is explained by differences between cultivars that remain constant over the two time points. For example, when cultivars that are susceptible early in the season (*t*_1_) remain susceptible late in the season (*t*_2_). Second, is the effect of pathogen. In this case, correlations may arise due to the highly local progression of epidemics characteristic of splash-dispersed pathogens or other cultivar-independent factors. For example, plots that had more disease early in the season would also have more disease late in the season. In other words, when considering the effect of pathogen, we investigate how well differences between plots (of the same cultivar, but located in separate lots) early in the season correlate with differences late in the season. These differences between plots may arise due to different amount of local inoculum or due to different local microenvironmental conditions. We note that genotype-environment interactions may contribute to both of these effects, but for simplicity we will refer to the two effects as “host effect” and “pathogen effect”. To separate the two effects, we performed a more detailed analysis of the correlation.

To determine the effect of host, instead of using two values per cultivar that correspond to two replicate plots (as we did in Fig. 4 of the main text), we used the average of these two values for each cultivar (by averaging over about 32 leaves belonging to each cultivar at each time point, with the exception of cultivar CH Claro that had about 670 leaves per time point). We then determined correlations between these average values in *t*_1_ and in *t*_2_. The resulting correlations are shown in Fig. E4.1 and in columns “cv” of the Table E4.1. The highest correlation is observed between *ρ*_lesion_ at *t*_1_ and *ρ*_leaf_ at *t*_2_ The best predictor for late-season PLACL is *ρ*_lesion_.

To determine the effect of pathogen, we normalized means over each plot in the following way. At each time point, we subtracted from each mean over plot (averaged over maximum 16 leaves) a grand mean over both plots (averaged over maximum 32 leaves) that belong to the same cultivar and time point. To balance the dominating effect of cultivar CH Claro, we used only two data points for it so that each point represents the difference between the mean over all 21 plots of CH Claro in one lot and the grand mean over 42 plots from both lots. The resulting correlations are shown in Fig. E4.2 and in columns “normed” in Table E4.1. The highest correlation is observed between PLACL at *t*_1_ and *ρ*_lesion_ at *t*_2_ The best predictor for late-season PLACL is *ρ*_lesion_.

Correlations corresponding to the effect of host with respect to the same measure (PLACL, *ρ*_lesion_, or *ρ*_leaf_) reveal the degree to which cultivars keep their ranking according to this particular measure between *t*_1_ and *t*_2_ (see diagonal panels in Fig. E4.1). In these cases, correlation coefficients quantify the degree of consistency between *t*_1_ and *t*_2_ of host resistance to PLACL (Fig. E4.1A), *ρ*_lesion_ (Fig. E4.1E) and *ρ*_leaf_ (Fig. E4.1H). As we see from Fig. E4.1A, PLACL at *t*_1_ has no significant correlation to PLACL at *t*_2_. Hence, we cannot predict differences in host damage between two cultivars late in the season based on differences in host damage early in the season. Such prediction is more likely to be achieved if we considered differences between one of the most susceptible and one of the most resistant cultivars. However, when we take into account all cultivars, on average there is no significant correlation. In contrast, the correlation between *ρ*_lesion_ at *t*_1_ and *ρ*_lesion_ at *t*_2_ is moderately strong and highly significant. We conclude that differences between cultivars according to *ρ*_lesion_ are more consistent than differences according to PLACL.

Correlations corresponding to the effect of pathogen reveal the degree to which plots of the same cultivar keep their ranking between *t*_1_ and *t*_2_. In contrast to the “host effect” described above, PLACL at *t*_1_ has a significant slightly negative correlation to PLACL at *t*_2_ with respect to “pathogen effect” (Fig. E4.2A). *ρ*_leaf_ at *t*_1_ and *ρ*_leaf_ at *t*_2_ exhibit a moderate correlation (Fig. E4.2I). Hence, differences between plots of the same cultivar are more consistent between *t*_1_ and *t*_2_ with respect to *ρ*_leaf_ than with respect to PLACL.

To determine their relative contributions to the overall correlations shown in Fig. 4 of the main text, we compare contributions of “host effect” (Fig. E4.1) and “pathogen effect” (Fig. E4.2). Correlation between PLACL at *t*_1_ and *ρ*_lesion_ at *t*_2_ due to “pathogen effect” is positive and significant (Fig. E4.2D), while the same correlation due to cultivar effect is not significant (Fig. E4.1D). Hence, we conclude that the overall positive relationship between PLACL *t*_1_ and *ρ*_lesion_ *t*_2_ (Fig. 4D of the main text) is caused mainly by “pathogen effect”. In contrast, there is no significant relationship between *ρ*_lesion_ at *t*_1_ and *ρ*_leaf_ at *t*_2_ due to pathogen effect (Fig. E4.2G), but there is a strong positive correlation due to “host effect” (Fig. E4.1G). This indicates that the overall positive correlation arises due to “host effect”. Interestingly, *ρ*_lesion_ exhibits a moderate positive correlation between *t*_1_ and *t*_2_ due to “cultivar effect” but a negative correlation due to “pathogen effect”. Reason for that phenomenon remains an open question.

Since cultivar CH Claro was replicated more than any other cultivar (a total of 42 plots), we analyzed it separately to see if the “pathogen effect” discussed above is visible in data from one cultivar. However, Spearman’s correlation test did not show any significant correlation with respect to means over each of 42 plots between *t*_1_ and *t*_2_ for any combination of the three quantities (PLACL, *ρ*_lesion_ and *ρ*_leaf_).

Thus, separating the “host effect” and “pathogen effect” allowed us to gain insight into the source of the correlations over time between different measures of host resistance and to better assess their predictive power. Although this analysis did not reveal extremely strong correlations, we expect, that disease forecasting models could benefit from having pathogen reproduction, not only host damage, as an explanatory variable. Combined with incidence measurements, the method presented allows for detailed quantification of pathogen population and its reproductive potential.

**Table E4.1:** Spearman’s correlations between measures of resistance in *t*_1_ and *t*_2_. Nonsignificant correlations in parentheses (*p >* 0.01). Correlations in columns “raw” are calculated using means of each plot for each time point, in columns “cv” using mean over all plots of a cultivar for each timepoint and in columns “normed” using difference of plot mean from cultivar mean for each time point as explained in the text.

| | PLACL($t_1$) | | | $\rho_{\text{lesion}}(t_1)$ | | | $\rho_{\text{leaf}}(t_1)$ | | |
| --- | --- | --- | --- | --- | --- | --- | --- | --- | --- |
|  | raw | cv | normed | raw | cv | normed | raw | cv | normed |
| PLACL($t_2$) | (-0.02) | (0.03) | -0.11 | 0.39 | 0.41 | 0.27 | 0.30 | 0.37 | (0.081) |
| $\rho_{\text{lesion}}(t_2)$ | 0.25 | (0.07) | 0.44 | 0.22 | 0.41 | -0.20 | 0.44 | 0.43 | 0.35 |
| $\rho_{\text{leaf}}(t_2)$ | 0.17 | (0.06) | 0.38 | 0.39 | 0.53 | (-0.062) | 0.50 | 0.52 | 0.39 |

**Figure E4.1:**
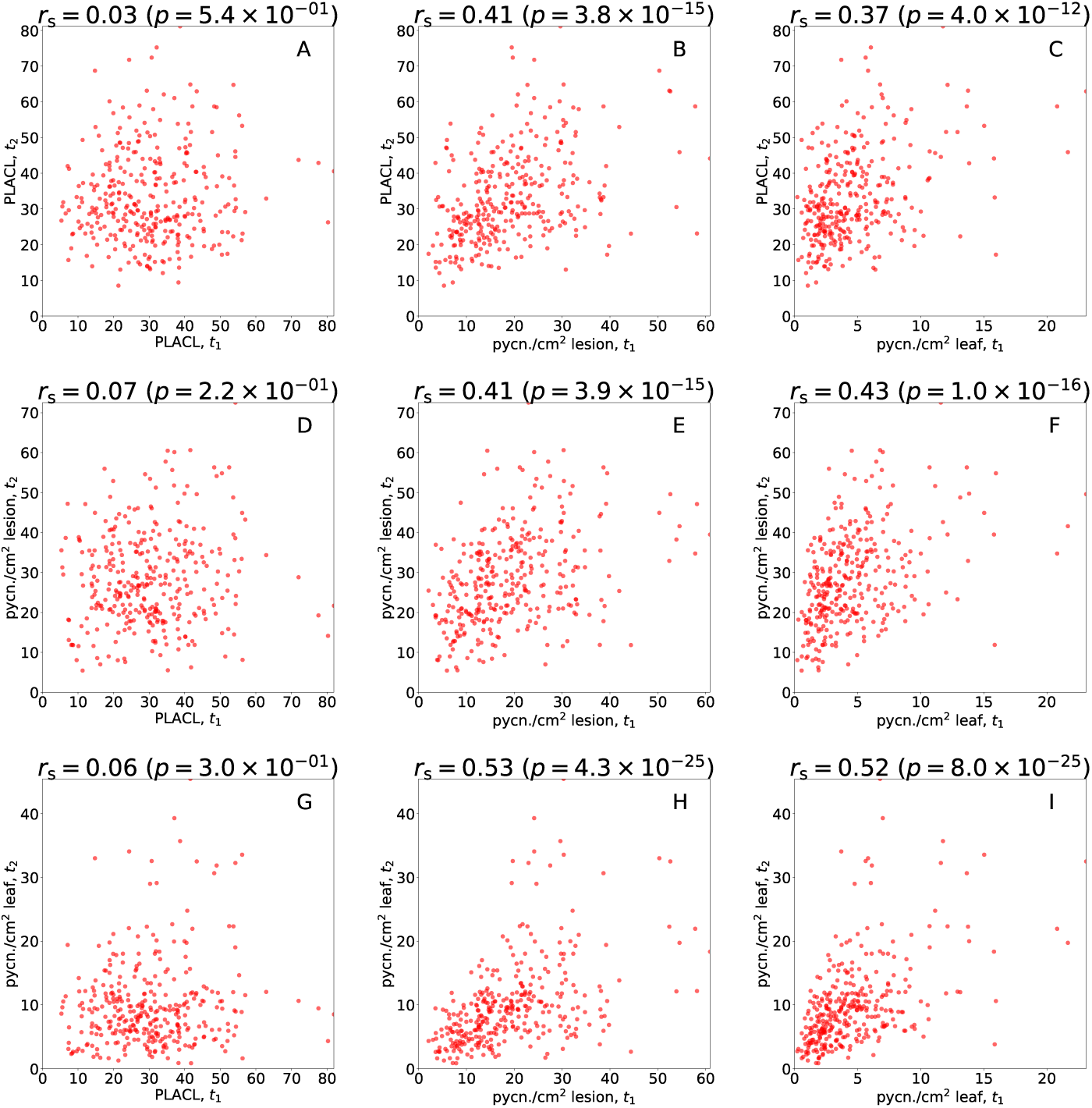
Correlation of measures of resistance (means over plots of the same cultivar) between the first time point (*t*_1_) and the second time point (*t*_2_). Each data point represents an average over about 30 leaves of one cultivar belonging to two replicates plots. Degree of correlation is quantified using Spearman’s correlation coefficient, *r*_s_.

**Figure E4.2:**
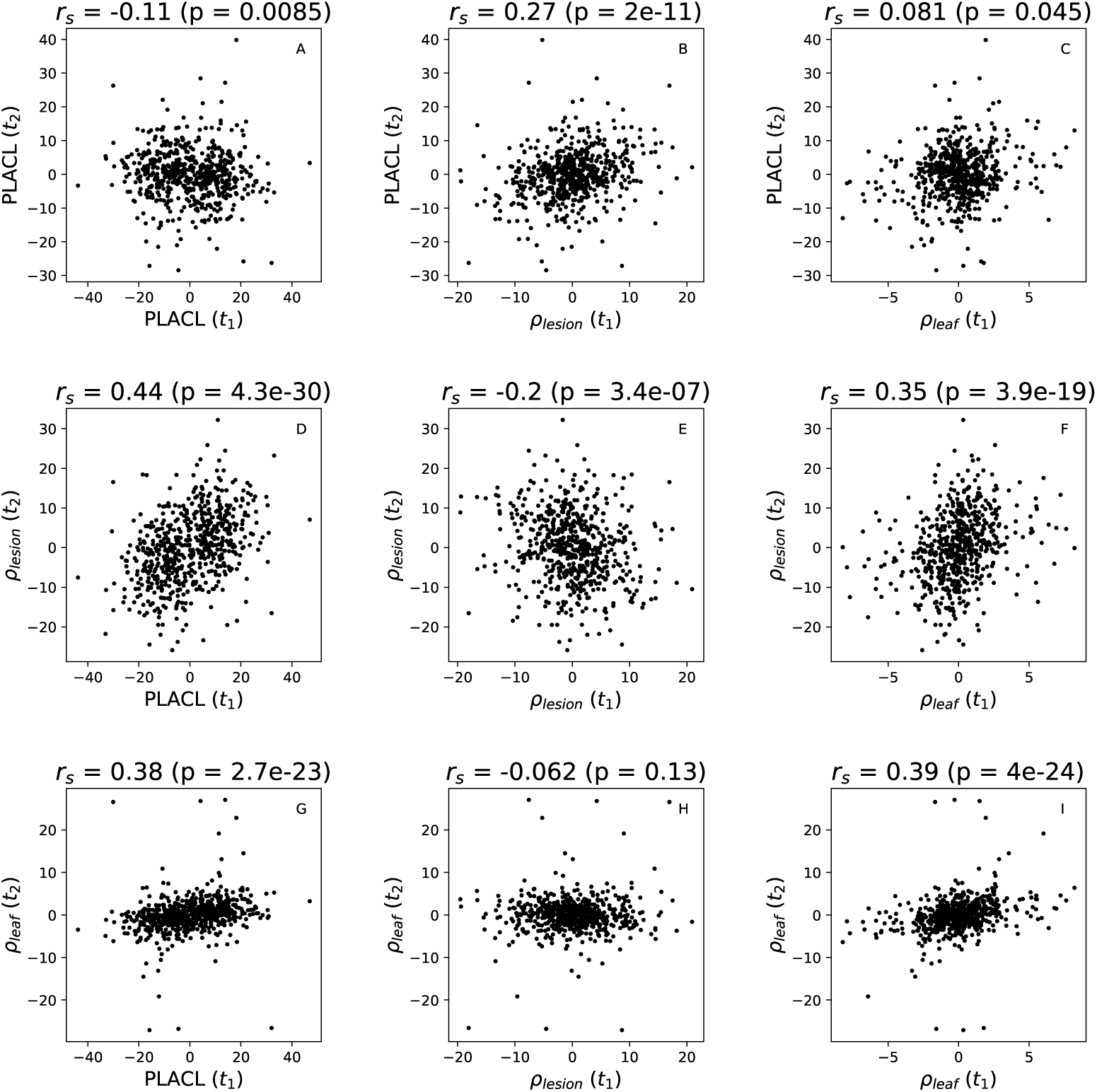
Correlation of measures of resistance between *t*_1_ and *t*_2_ within cultivars. Data from each cultivar is represented by two data points. For each point, the *x*-value represents the difference between the mean over an individual plot and the grand mean over two plots at *t*_1_ and the *y*-value is the same difference taken at *t*_2_. Each data point represents an average over about 15 leaves of one cultivar belonging an individual plot. The degree of correlation is quantified using Spearman’s correlation coefficient, *r*_s_.

